# Feedback between stochastic gene networks and population dynamics enables cellular decision-making

**DOI:** 10.1101/2023.11.21.568034

**Authors:** Paul Piho, Philipp Thomas

## Abstract

Phenotypic selection occurs when genetically identical cells are subject to different reproductive abilities due to cellular noise. Such noise arises from fluctuations in reactions synthesising proteins and plays a crucial role in how cells make decisions and respond to stress or drugs. We propose a general stochastic agent-based model for growing populations capturing the feedback between gene expression and cell division dynamics. We devise a finite state projection approach to analyse gene expression and division distributions and infer selection from single-cell data in mother machines and lineage trees. We use the theory to quantify selection in multi-stable gene expression networks and elucidate that the trade-off between phenotypic switching and selection enables robust decision-making essential for synthetic circuits and developmental lineage decisions. Using live-cell data, we demonstrate that combining theory and inference provides quantitative insights into bet-hedging-like response to DNA damage and adaptation during antibiotic exposure in *Escherichia coli*.

## INTRODUCTION

Cells make decisions in response to changes in gene expression, which is surprisingly noisy even among genetically identical cells facing the same environmental conditions [1–3]. Such cellular noise arises from randomness in the biochemical reactions synthesising proteins. Since these proteins are involved in gene regulatory networks, stochasticity in expression levels can impact cellular functions, cell proliferation, and survival. The interactions between gene networks and population dynamics give rise to phenotypic selection even in clonal populations [3–5]. Phenotypic selection can have functional consequences in development [6, 7], how cells respond to stress [8–10], and drug resistance [11–13]. Understanding these consequences is important to enhance the function of synthetic circuits inside cells [14–16].

The Gillespie algorithm is widely used to simulate stochastic gene expression [1]. The method exactly simulates reaction dynamics at cellular scales, but it implicitly assumes that cells are static and gene expression occurs in isolation from dynamic cellular context. Recent studies have challenged this static view of cells by examining the effect cell division has on gene expression through mechanisms of partitioning of molecules [17–20]. Some studies showed that when cells compete for growth, the distribution across growing populations differs from isolated lineages as observed in the mother machine [21–23]. However the differences between such measures are not well understood when gene expression affects cell division.

It is becoming increasingly clear that gene expression noise contributes to cell-to-cell variation in cell growth [24, 25] and cellular timings [26]. A range of studies have focused on modelling interdivision time through the expression of a division protein hitting a set target level from a fixed basal level [27–31]. The model recovers correlations in interdivision time and cell size compatible with the adder division rule in bacteria. However, a single division protein consistent with time-lapse observations cannot always be identified, and several cues may contribute to phenotypic selection when cells respond to stress or drugs.

More generally, the coupling of gene expression and cell division involves feedback, where gene expression affects cell division frequency, which modulates expression levels. For example, competition between cells decreases the interdivision time and in turn increases the frequency with which molecules are partitioned at cell division [23]. When gene expression affects the interdivision time [32, 33] – which we refer here to as division-rate selection – it leads to additional competition. Selection due to differences in division rate is thus distinguished from natural selection arising from a competitive advantage of fast-growing subpopulations.

Natural selection can, in principle, be probed through switching off competition. This can be achieved, for example, by culturing cells in a mother machine [34]. Yet differences in division rates are expected to impact expression levels even in isolated cells and can also be engineered artificially [14]. A quantitative theory of the interactions between gene networks and population dynamics is still missing and it thus remains elusive how to distinguish natural selection from selection on division rate.

We propose a stochastic framework to model cells as agents that divide in response to intracellular stochastic reaction networks. The model allows us to probe how gene expression contributes to cell proliferation and natural selection and how these effects shape gene expression. Specifically, we show that the division distributions in growing cell populations differ from the first passage distributions of conventional stochastic reaction networks due to the consequences of division rate, natural selection, and cell history. We provide accurate but tractable approximations that provide insights into wide ranges of parameter space and allow us to quantify selection effects from time-lapse observation data.

## RESULTS

### Analytical framework for stochastic agent-based modelling of selection in gene networks

We developed a stochastic agent-based formulation of a growing clonal cell population that couples the internal stochastic gene regulatory dynamics and the population dynamics via cell division. In the model, each cell is represented by an agent (Fig. 1A) that contains a gene regulatory network composed of biochemical species 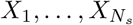 that react through *R* intracellular reactions. Consequently, the state of each cell is given by its age *τ* and the molecular content ***x***. The cells divide at an ***x*** and *τ* -dependent rate *γ*(***x***, *τ*) where the dependence of *γ*(***x***, *τ*) on ***x*** encodes the effects of selection on ***x***. For example, if *γ*(***x***, *τ*) is a monotonically increasing function in ***x*** we have positive selection, and negative selection in the case of monotonically decreasing dependence. There is no selection on ***x*** if the division rate depends only on *τ*. Such agent-based models of cell populations can be exactly simulated by the extended first division algorithm (see Materials & Methods).

**FIG. 1.**
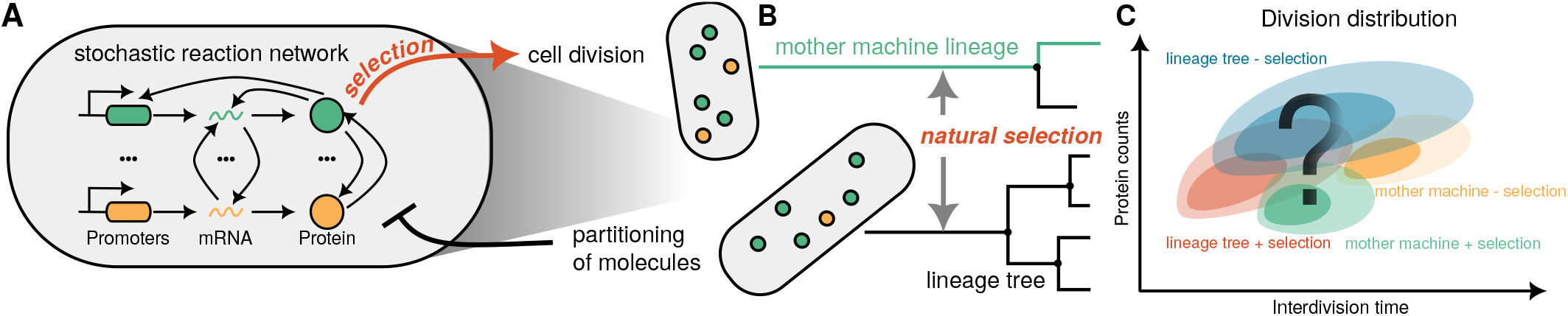
Quantifying interactions between gene networks and population dynamics with agent-based modelling of clonal populations. **(A)** Cartoon of the agent-based model where intracellular reaction network couple to cell division. Intracellular reactions affect cell division rate (red arrow) while partitioning of molecules dilutes cellular expression levels (black repressive arrow). **(B)** Lineage statistics of agent-based models measure distributions across lineage tree resulting from competition of cells through natural selection. A mother machine lineage (green line) follows a single cell in the lineage tree starting from an ancestral cell and following each daughter cell with equal probability. Mother machine sampling avoids natural selection, i.e. competition of cells for growth (red arrow). **(C)** Illustration of division distributions for a cell to divide at a given age and protein count for a lineage tree of cells including division-rate selection and natural selection (red), a lineage tree without division-rate selection (blue), and the corresponding mother machine lineages (green and orange).

Agent-based stochastic simulations are time- and resource-consuming, especially in the context of parameter inference, when large numbers of parametrisations need to be checked. We can make analytical progress by considering the long-term behaviour of the mean number of cells *n*(***x***, *τ, t*) with molecule counts ***x*** and age *τ* that grows exponentially with time *t* (SM Section S1). Normalising this quantity leads to a stable snapshot distribution 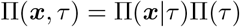, where Π(***x***|*τ*) is the (conditional) snapshot distribution of molecule numbers for cells with a given age *τ*, and Π(*τ*) is the age distribution given by 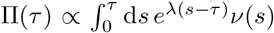. The age distribution is consistent with the age-structured models of McKendrick and von Foerster [35]. In their models, the interdivision time distribution *ν* affecting cell age and division and determining the population growth rate *λ* is typically assumed to be fixed parameters or provided by experimental data. However, the (conditional) snapshot distribution is less studied and, as we will see, interdivision time distribution generally depends on the gene network dynamics in the presence of selection.

We derived the first passage distribution *ν*(***x***, *τ*) for a cell to divide at state ***x*** and age *τ*, henceforth called the division distribution for brevity:

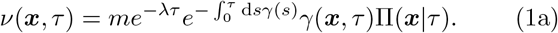

Here, *m* is the number of offspring at cell division and *γ*(*τ*) = 𝔼_Π_ [*γ*(***x***, *τ*)|*τ* ] is the marginal division rate with respect to the conditional distribution Π(***x***|*τ*). The snapshot distribution Π(***x***|*τ*) of gene expression satisfies a master equation

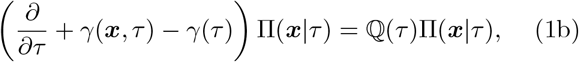

which has to be solved along with a boundary condition that connects the molecule numbers at cell birth and division:

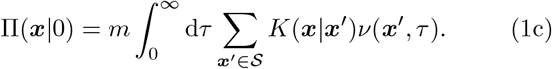

Biochemical reactions are encoded by the transition matrix

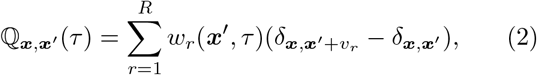

where *δ* is the Kronecker delta, *w*_*r*_(***x***, *τ*) are the (potentially cell cycle-dependent) reaction propensities, and *v*_*r*_ is the reaction stoichiometry. The partitioning kernel 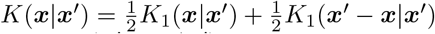 where *K*_1_(***x***|***x***′) and *K*_1_(***x***′ − ***x***| ***x***′) are the marginal distributions of molecules inherited by the two daughter cells from the mother cell with intracellular state ***x***′.

The theory highlights the intricate feedback that exists between gene expression and cell division and underlies division-rate selection. The division rate multiplies the gene expression distribution in the division distribution (1a), meaning that cells where *γ*(***x***, *τ*) is low are underrepresented while cells with high division rate are overrepresented. The division distribution then determines the distribution at cell birth (1c) and the time-evolution of the gene expression distribution (1b). The latter differs from chemical master equation models as the frequency of gene expression phenotypes is not only determined by biochemical reactions but modulated by division-rate selection throughout the cell cycle.

Natural selection contributes to this feedback through the exponential dependence of the division distribution on the population growth rate. Cells dividing slower than the population doubling time are under-represented in the division distribution compared to cells that divide faster. The strength of natural selection is controlled through the number of offspring at cell division. For example, for a lineage tree of cellular agents, we have *m* = 2 and the population growth rate *λ* needs to be computed self-consistently through normalising (1a) while in the mother machine setting, i.e. considering an isolated cell lineage, we have *m* = 1 and *λ* = 0. The difference between mother machine lineages and lineage trees provides a measure of natural selection [36–39]. The theory extends to cell growth and cell size control through an effective division rate (see Materials & Methods) and thus provides a coarse-grained view of gene expression-induced selection effects.

### Division-rate vs natural selection in the telegraph gene expression model

To study selection effects, we consider the telegraph model (Fig. 2A) where a promoter switches between active and inactive states. Once the promoter is activated, a protein is produced in random bursts. Selection in our model is introduced through the division rate, which we assume to be of the form:

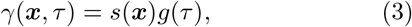

where *s*(***x***) describes the effects of gene expression levels ***x*** on the division rate and *g*(*τ*) is the division rate in the absence of selection on the gene expression (see also Materials & Methods on modelling growth rate-dependent selection). Our main equations (Eqs. 1a to 1c) cannot be solved in closed form because they are essentially an infinite system of coupled integro-ODEs. An analytical solution is only known in the special case with deterministic divisions and without selection [20, 40, 41].

**FIG. 2.**
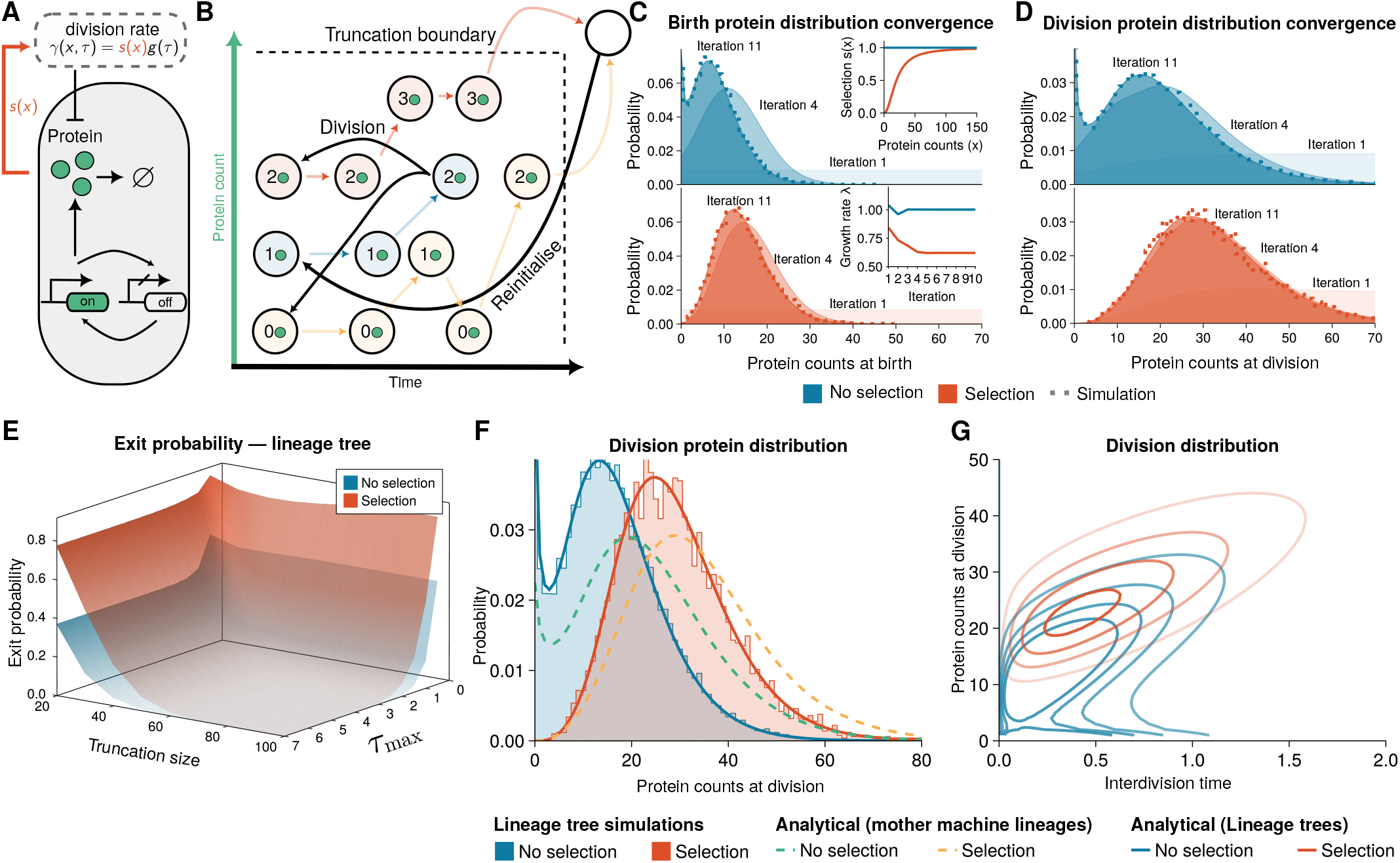
Finite state projection (FSP) enables accurate prediction of selection on gene expression noise. Schematic illustration of the agent-based model. Intracellular dynamics of stochastic gene expression are modelled by the telegraph model for bursty expression of a protein. The protein is binomially partitioned at cell division. The selection effect is introduced through a division rate increasing with expression (details in SM Section S3 A). **(B)** Illustration of the FSP algorithm. Intuitively, gene expression (light arrows) drives cells to either commit to division or cross the truncation boundary, leading to their states being reinitialised (dark arrows). **(C-D)** Birth and division distributions for the model converge in a few iterations and agree with agent-based simulation of lineage trees obtained using the first division algorithm (Materials & Methods, see SM Fig. S1 for mother machine lineages). Top inset of **C** shows the used selection functions (*s*(***x***) = 1 for no selection, *s*(***x***) = 1/((20/*x*)^2^ + 1) for selection on the gene expression). Bottom inset shows the convergence of the population growth rate *λ* computed in Method. **(E)** The exit probability due to FSP truncation decreases with truncation size and time horizon *τ*_max_. **(F)** Summary distributions without (blue solid line) and with selection (red solid line) are compared to direct simulations of the lineage trees (shaded areas). Our method is in agreement with the simulation results. Distributions obtained using mother machine lineages (dashed) are also shown. **(G)** FSP solution of the agent-based model predicts bimodal division distribution without division-rate selection (blue lines correspond to levels of equal probability) and an unimodal division distribution in the presence of division-rate selection.

To overcome this challenge, we developed a powerful approximation based on the finite state projection (FSP) method (see Fig. 2B, Materials & Methods). In brief, the method consists of restricting the dynamics to a finite subset of the state space 𝒳 ⊂ 𝒮 for ***x*** and solving the dynamics of Eqs. 1a to 1c up to a finite cell age *τ*_max_. This is akin to what is done in the standard FSP method [42] but, in addition, we introduce (i) cell division events that split the dividing cell via partitioning of molecules leading to *m* newborn cells and (ii) events that reinitialise single-cell trajectories once they leave the prescribed state space (Fig. 2B).

For the case without selection, we set the expression-dependent selection function in the telegraph model to be *s*(***x***) = 1 and a Hill function for positive selection (Fig. 2C, top inset). We computed the birth and division distributions for a fixed truncation size (Fig. 2C,D) along with the corresponding growth rate (Fig. 2C, bottom inset). We observe that the iterative scheme converges (Fig. 2C,D) and the method allows us to compute the exit probability that arises from state space- and cell age truncation ((M1), Fig. 2E). Equation (M5) corresponds to an Euler-Lotka equation of population dynamics [43] up to an error term that involves the exit probability. The exit probability measures the proportion of cells that reach the truncation boundary and are thus reinitialised instead of dividing. It decreases monotonically with maximum age *τ*_max_ and truncation size (Fig. 2E) guiding the accuracy of the approximation. The resulting FSP solution (Fig. 2C-D shaded area) also agrees well with simulations using the First Division Algorithm (dots, see Materials & Methods).

Interestingly, our analysis predicts bimodal distributions without selection on the gene expression while the distributions with selection are unimodal (Fig. 2F-G). Bimodality in the absence of selection arises from slow promoter switching [44]. Such long lived transcriptional states can arise from transcription factor-mediated looping of DNA as observed in the lactose operon of *Escherichia coli* [45]. To clarify whether the absence of the zero-mode is due to the slow-growing sub-population being outcompeted or due to division-rate selection on the gene expression, we also computed the division distributions of mother machine lineages (Fig. 2F). We observe qualitatively similar division protein distributions for both population lineage trees and mother machine lineages, and division-rate with natural selection in the population skewing distributions towards lower expression levels. Intuitively, this can be understood through Equation (1a), which is proportional to the division rate while division times longer than the population doubling time are exponentially suppressed. Importantly, division-rate selection on the gene expression accounts for the absence of the slow subpopulation in both measures. This highlights that selection in the telegraph model is primarily driven by division rate.

### Growth feedback reinforces lineage decisions in multi-stable gene regulatory networks

We wondered whether simple patterns of selection are important for cell fate decisions. Traditionally, cell differentiation is associated with multi-stability in gene regulatory networks shaping Waddington landscapes [46]. It is known that protein expression modelled by the birth-death process leads to unimodal distributions both in mother machine and lineage trees [22]. We therefore considered positive feedback loops that are commonly associated with bistability and bimodal distributions (Fig. 3A). In the model without selection, a protein is expressed from a promoter which in turn increases its own expression. The production rate is modelled as constant basal transcription rate *α* along with a Hill-type function creating a positive feedback loop. Simulation of population histories (a random lineage of a lineage tree [36]) and a mother cell lineage show large fluctuations (Fig. 3B) leading to long-tailed or bimodal distributions (Fig. 3C,D).

**FIG. 3.**
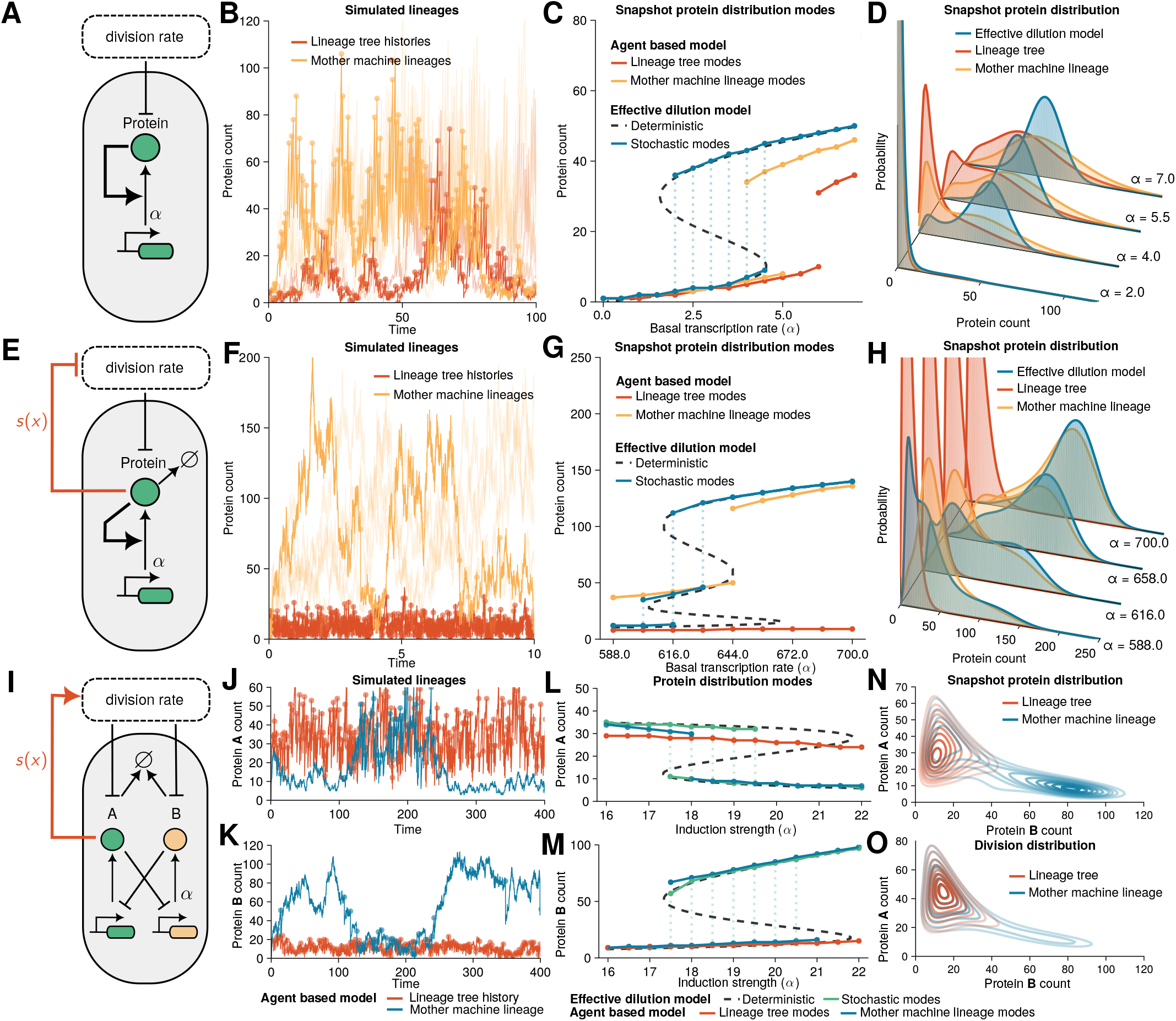
Selection reinforces lineage decisions in cellular switches. **(A-D)** Agent-based model of transcriptional feedback. **(A)** Illustration of the transcriptional feedback model where protein expression promotes its production with parameter *α* corresponding to a basal transcription rate. Proteins are partitioned binomially at cell division. **(B)** Agent-based simulations of lineage tree histories and mother machine lineages (*α* = 4.0) obtained using the First Division Algorithm (see Materials & Methods) display switching between low and high protein expression states. Individual cell divisions are shown as dots to indicate the time-scale of cell divisions. **(C)** Modes (local maxima) of the snapshot protein distributions. FSP solutions of the agent-based model predict bimodal distributions of mother machine (orange) and lineage trees (red) over intermediate basal production rates. The EDM (black and blue) agrees well with the mother machine solution. **(D)** FSP solutions show a transition from unimodal to bimodal distributions. **(E-H)** Agent-based model with transcriptional and growth feedback (*s*(***x***) = *k*_2_/((*x/K*_2_)^4^ + 1)) where protein synthesis inhibits division rate and promotes its own production rate. EDM and mother machine lineages have three stable modes while lineage trees show that fast-dividing cell lineages take over the population. **(F)** Agent-based simulations of mother machine lineages (basal transcription rate *α* = 644.0) show switching between low and high protein levels while in lineage tree histories the fast dividing lineages determine the cell fate. **(I-O)** Agent-based model of the genetic toggle switch. Induction strength *α* corresponds to the maximal transcription rate of the protein B. The protein A is under selection via selection function of the form *s*(***x***) = *k*_3_/((*K*_3_/*x*_*A*_)^2^ + 1). **(N-O)** Snapshot and division protein distribution display multimodality and a long tail, respectively, in the mother machine lineages (blue, *α* = 18.9). The fast-dividing subpopulation is selected in the lineage trees (red). Parameters and model details in SM Sections S3 B and S3 C.

This phenomenon can be understood in terms of slow switching between discrete states of an effective dilution model (EDM). These effective reaction networks add one dilution reaction per species to the reaction network and can be simulated easily using ODEs or Gillespie simulations. Our analysis of the ODE model of the network reveals the S-shaped multistable response to changes in the basal transcription rate (Fig. 3C, dashed black line).

Characteristically, traces simulated through the Gillespie simulations switch between two equilibria of high and low gene expression that correspond to modes in the stationary probability distributions (Fig. 3C, solid blue line). Both the lineage tree and mother machine lineages show bimodal snapshot (Fig. 3C) and division distributions (SM Fig. S2) and thus qualitatively agree with the EDM over large parameter regimes. However, a quantitative comparison of the snapshot protein distributions reveals different noise characteristics. EDM gives a better approximation to the mother machine lineage formulation than the lineage trees but underestimates the noise since it provides only an effective description of cell divisions (Fig. 3D).

Due to the presence of random switching, differentiation of this circuit can only be achieved when an external parameter is varied. Yet, how to design circuits that differentiate irreversibly in the absence of fine-tuned signals remains unclear. To this end, we study an extended circuit where additional feedback is introduced to the model via division rate (Fig. 3E). Cell division is modulated through the repressive Hill-type selection function. The extra feedback changes the parameter regime where multimodality appears and in some parameter regimes introduces an additional steady state in the EDM (Fig. 3G).

Simulations of mother machine lineages show switching between the tree expression levels with very slow cell divisions in the highly expressed states (Fig. 3F, orange line). In stark contrast, the histories of population lineage trees display fast divisions selecting only the low-expression state (Fig. 3F, red line) due to the negative feedback between expression and division rate. The higher expression level states in a population get rapidly outcompeted by fast-dividing cells. Comparing the distribution modes of the different measures, we observe that the EDM model explores all three states (Fig. 3G-H, SM Fig. S3). The mother machine lineages mostly explore the two high expression states corresponding to slow-dividing cells while the population lineage trees display exclusively the lowly expressed state that promotes fast cell proliferation. A similar effect, with the fast-dividing subpopulation being selected in lineage trees, is observed when removing the transcriptional feedback loop, which also includes an additional steady state due to growth feedback (SM Fig. S4).

We asked if selection could provide a general mechanism for lineage decisions beyond the single gene feedback models. We considered the genetic toggle switch as a common motif in synthetic biology, which comprises two antagonistic proteins inhibiting each other (Fig. 3I). The protein *A* is assumed to be under selection through the Hill-type selection function. The circuit dynamics are non-trivial as selection only acts on protein *A* while the dynamics are tightly coupled with protein *B*.

To analyse the balance between the division-rate selection and natural selection, we again compared the mother machine lineages with lineage trees. The trajectories of the mother machine lineages explore fast and slow dividing states while lineage tree histories settle into the fast dividing state where protein *A* is more highly expressed (Fig. 3J,K). The effects are confirmed by estimating the modes of the distribution computed using FSP where mother machine lineages are bimodal in close agreement with the EDM (Fig. 3L,M). In contrast, the protein distributions in lineage trees are unimodal (Fig. 3L-N). Interestingly, we observe that the bimodality in protein distributions of mother machine lineages was weaker at division (Fig. 3N) than in snapshots (Fig. 3O), implying that division-rate selection shapes the balance between these states.

The effect of division-rate selection in mother machine lineages can be seen as an over-representation of slow-dividing cells in snapshots compared to the division distributions that occur because cells spend more time in these slow states. This suggests that division-rate selection acts on division times in mother machine lineages in similar directions to natural selection on lineage trees. However, only due to natural selection, do fast-dividing cells eventually outcompete the slow-dividing ones (SM Fig. S6, S7). The coupling between gene expression and growth thus represents a robust strategy to implement cell fate decisions in natural and synthetic populations.

### Model-based inference predicts DNA damage response from division-rate selection

We now apply our modelling framework to understand how cells respond to stress. To this end, we investigate the activity of the SOS pathway quantified using the SOS promoter *PsulA* driving the expression of a fluorescent reporter [47]. Surprisingly, we found that the expression of the reporter is highly heterogeneous even in unstressed conditions (Fig. 4C, shaded blue bars). We use a bursty gene expression model to quantify the expression of *PsulA* (Fig. 4A, SM Section S4 B). Although we only model a single gene, burstiness describes both expression noise and upstream variability [48]. The simple model fits the data well (blue line, Fig. 4B-C).

**FIG. 4.**
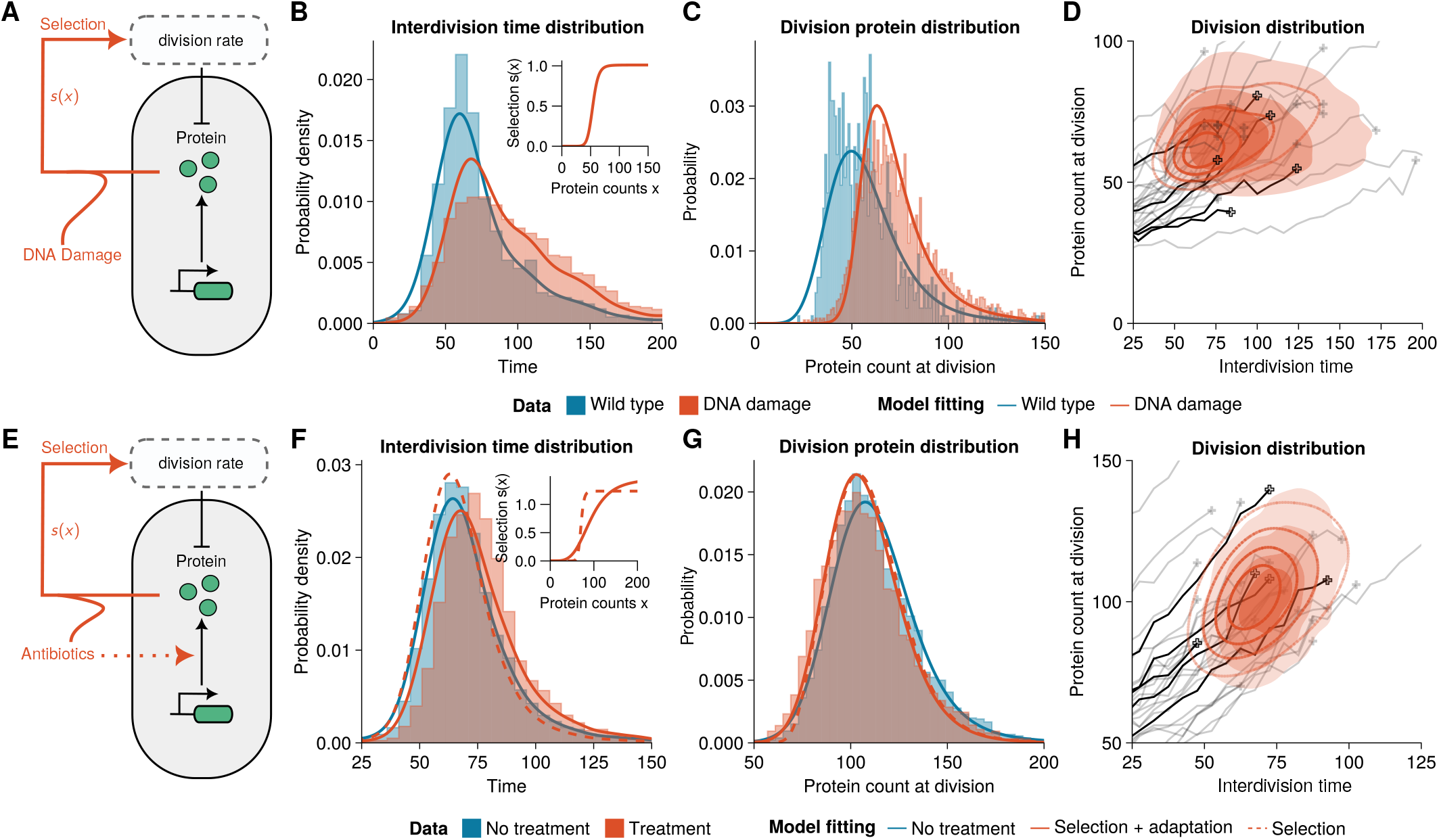
Inference of selection effects in *Escherichia coli* cells. **(A)** Agent-based model of the SOS response involves a bursty gene expression model with an age-dependent division rate and binomial partitioning at cell division. DNA damage is induced through a gene expression-dependent modulation of the division rate. **(B-C)** Interdivision time and protein distributions from the mother machine lineages (fluorescent reporter, medium growth conditions [47]) in unstressed (blue area) and damage-induced mother machine lineages (red area) are well fit by the agent-based model (*m* = 1, see SM Section S4 B for fit parameters). The inset shows the selection function *s*(***x***) obtained in damage-induced conditions. **(D)** The division distribution of the fitted model shows a distinctly peaked distribution (red lines) that compares well with the experimental distribution (red-shaded area) and single-cell traces of the data (representative traces shown in black and grey). **(E)** Agent-based model of antibiotic resistance gene expression using a bursty gene expression model with age-dependent division rate. Antibiotic treatment response involves positive feedback of protein expression and division through a gene expression-dependent division rate and adaption of gene expression parameters. **(F-G)** Statistics of the division distribution of the lineage trees (fluorescent fusion-protein genome-integrated reporter, [36]) in untreated (blue area) and antibiotic-treated cells (red area). Lineage tree data is well fitted by the agent-based model combining drug-dependent selection and adaptation (solid red line), but not using selection alone (dashed red line, *m* = 2, parameters in SM Section S4 B). The inset shows the corresponding selection functions *s*(***x***) obtained in treated conditions. **(H)** The division distribution under treatment shows a peaked distribution (red lines) that compares well with the experimental distribution and single-cell traces of the data (representative traces shown in black and grey).

In the model, we assumed the division rate to be in the form of Eq. (3), where *s*(***x***) models a damage-induced modulation and *g*(*τ*) is the division rate in unstressed conditions. The latter can be inferred directly from the interdivision time data of the wild type (*s*(***x***) = 1) using a non-parametric kernel density estimator (Fig. 4B, shaded blue bars) and fits well the interdivision time distributions. To infer the reaction kinetics in unstressed conditions, we convert fluorescence measurements to molecule numbers (SM Fig. S8) and define a likelihood function via the division distribution *ν*(***x***, *τ*) that can be maximised using Bayesian optimisation (see Materials & Methods, [49]) to fit the reaction parameters, i.e. burst size and burst frequency.

When DNA damage was induced, cells displayed a long tail of slowly dividing cells and increased levels of SOS pathway expression (Fig. 4B,C shaded red). Intuitively, this could suggest not only up-regulation of the SOS pathway as a stress response but also positive selection on SOS levels. We investigated this by inferring stress-induced selection function *s*(***x***) (Fig. 4B inset, see SM Section S4 B), but kept the reaction parameters as in unstressed conditions, which provided good agreement with the data (Fig. 4B,C solid red line). We hence conclude that upregulation of SOS expression is not required to fit the data. We wondered whether these effects could equally be explained through a neutral model without selection but with different interdivision time distributions. The latter gave a worse fit as it overestimated the protein-interdivision time correlation (SM Fig. S9).

Our findings thus suggest that the DNA damage response is driven mainly by division-rate selection in this system. The observation alludes to a possible bet-hedging strategy in *Escherichia coli* where SOS expression is heterogeneous in unstressed conditions to better deal with environmental changes in stress levels [50].

### Antibiotic resistance gene expression involves division-rate selection and adaptation

As a second application, we considered time-lapse data of *Escherichia coli* expressing an antibiotic resistance gene *SmR*, an efflux pump conferring resistance to *streptomycin* [36]. Again, we find that protein distributions are highly heterogeneous even in the absence of antibiotics and can also be fitted by a bursty model involving a single gene (Fig. 4E-G solid blue) parameterised through burst size and burst frequency.

We wondered whether antibiotic resistance gene expression represents a possible bet-hedging strategy similar to one observed for the SOS response (Fig. 4A-D). To this end, we again modelled the division rate via (3), where now *g*(*τ*) is the division rate in the absence of antibiotics (*s*(***x***) = 1) and *s*(***x***) is a drug-induced modulation of division rate. We converted fluorescence to molecule numbers (SM Fig. S10) and used Bayesian optimisation to fit the protein-dependent modulation of the division rate *s*(***x***) underlying the interdivision time and protein distributions under conditions where cells have been exposed to sub-lethal doses of antibiotics. While this division rate described well the reduction in protein expression (Fig. 4G dashed red), it failed to account for the increase in interdivision time (Fig. 4F, dashed red). Similarly, a model with no selection but matched inter-division time distributions did not fit the protein expression data (SM Fig. S11). Intuitively, the data suggested attenuated protein expression while cells divided slightly slower (Fig. 4F,G red area), which is inconsistent with the division-rate selection model.

We thus hypothesised that cells adapt gene expression during treatment. Indeed, allowing for selection on the protein as well as adjusting the rate parameters, modelling the adaptation to the new conditions, provides good agreement with the experimental data. The model fits the interdivision time (solid red in Fig. 4F), division protein (solid red in Fig. 4G) as well as the joint division distribution of protein count and interdivision time (solid red in Fig. 4H). Our results suggest that burst size roughly doubles after treatment (SM Section S4 B). This highlights that adaptation plays an important role in the drug response in *Escherichia coli*.

## DISCUSSION

We developed an agent-based framework for quantifying the feedback between stochastic gene expression, cell division and population dynamics. Our approach allows us to derive the division distributions of lineage trees directly accessible in live-cell imaging of growing cell populations. These distributions are generally distinct from the first passage distributions of the conventional chemical master equation due to the effects of division rate, natural selection, and cell history. Our findings thus provide a quantitative understanding of how natural selection and division rates shape phenotypes and allow decision-making in response to gene expression.

Our method enables accurate solutions of agent-based models and efficient parameter inference. Previous approaches typically relied on costly agent-based simulations [51]. To this end, we proposed a finite state projection solution. We identified the exit probability of crossing the state space, which serves as an error gauge and quality assurance of the numerical scheme resulting from truncating the state space and setting a finite maximum age for cells. However, the FSP method (like the master equation) presents a large system of coupled differential equations. Tensor-train methods and clever state space truncations have alleviated some of these challenges [52].

Multi-stable gene regulatory networks are fundamental in development, triggering cellular responses, and cell fate decisions. Naturally, these networks require environmental signals for robust decision-making, while otherwise, they are subject to continuing fluctuations that can lead to reversible phenotypic switching. Our methodology allowed us to understand several models implementing positive feedback between protein expression and division rate. We showed that positive feedback coupled with population dynamics leads to phenotypic switching of mother machine lineages but to phenotypic selection in the histories of lineage trees, demonstrating that mother machine lineages generally do not represent typical cell lineages. Intuitively, the mother machine samples lineages with a probability that decreases exponentially with the number of divisions, thus oversampling slowly dividing cells compared to lineage trees [36]. Importantly, division-rate selection presents a robust mechanism of irreversible lineage choice in a population. This mechanism requires slow-cycling progenitor cells as observed in stem cell lineages [7, 53, 54]. Mother machine lineages and lineage tree histories, which originate from the same ancestral cell, both display phenotypic switching initially but then gradually diverge over time due to natural selection (SM Fig. S6, S7). This trade-off could present a strategy for adaptation to changing environments.

The present framework provides multiple advantages over previously proposed stochastic modelling approaches for single-cell data. Importantly, our framework revealed irreversible cell fate decisions that cannot be captured using traditional chemical master equation models. The agent-based approach goes beyond the snapshot statistics because it explicitly accounts for age structure. This allows us to assess lineage statistics and dynamics, such as cell birth, division events, or certain cell cycle stages. As we have demonstrated, this information is readily available from time-lapse data and allows for parameter fitting and model selection. Although we focused on division rate, growth rate is coupled to gene expression and hence is an important factor for selection [9, 25, 55]. Our framework extends to cell size control through an effective division rate (see Materials & Methods) where selection strength is proportional to the cell growth rate. Analysing growth and cell size dynamics could disentangle the effects of division and growth rates, and provide further data insights [56].

Our agent-based theory provides crucial insights into how cells respond to stress and drugs based on gene expression and division feedback. To this end, we employed Bayesian optimisation for parameter inference on two single-cell datasets in *Escherichia coli* [36, 47]. The framework allowed us to efficiently integrate data from multiple conditions to identify selection effects and evaluate competing model hypotheses. Our stochastic model suggested the increase in SOS pathway expression in response to induced DNA damage [47] does not require upregulation of SOS expression but is achieved effectively through division-rate selection. This suggests that heterogeneous SOS expression in unstressed conditions could provide a bet-hedging strategy. A similar model applied to the response of an antibiotic resistance gene revealed that division-rate selection alone was not sufficient to explain the data but that cells adapt their gene expression during treatment.

Crucially, our analysis quantified changes in gene expression and interdivision times between different experimental conditions. Yet, we cannot exclude that selection effects are present even in supposedly neutral conditions. It remains to be seen whether differences between division-rate and natural selection can be quantified from time-lapse observations of individual lineage trees. For simplicity, we assumed a multiplicative selection model, Equation (3), where gene expression-dependent effects act independently from other sources effectively modelled through cell age. Studying the effects of cell-cycle-dependent feedback on selection would be an interesting avenue for future research.

Our new theory enables us to understand how variability in gene expression shapes phenotypes and propagates to population dynamics. We used the method to uncover mechanisms of cell fate decision-making, study how cells respond to stress, and adapt their gene expression programmes in response to drugs. Our findings thus substantially advance the understanding of how cells make decisions through coupling gene networks with growth and division. Looking beyond gene expression, our framework could more widely be used to study phenotypic selection on other processes that affect growth, such as mitochondrial turnover [57]. We expect our methods to provide insights into the role of heterogeneity in growth-associated diseases and drug resistance, such as in cancer [58].

## AUTHOR CONTRIBUTIONS

Conceptualisation: PP and PT. Methodology: PP and PT. Simulation and data analysis PP. Writing: PP and PT.

## CODE AVAILABILITY

All data needed to evaluate the conclusions in the paper are present in the paper and/or the Supplementary Materials. Codes for the First Division Algorithm and FSP-based solution of the agent-based models are available as a Julia package at https://github.com/pthomaslab/AgentBasedFSP.jl (permanent snapshot at https://zenodo.org/doi/10.5281/zenodo.10797749). Examples used in the paper can be reproduced using codes at https://github.com/pthomaslab/SelectionPaperExamples (permanent snapshot at https://zenodo.org/doi/10.5281/zenodo.10797757).

## MATERIALS & METHODS

### First Division Algorithm for exact agent-based simulation

We present an exact simulation algorithm to the agent-based model of cells in a growing population from time *t*_0_ to *T*. This is an extended version of the First Division Algorithm [22] and the Extrande thinning method for sampling division times [59]. The algorithm uses a lookup horizon ∆*t* over which the division rate can be bounded.

1. *Initialisation* Initialise. the cell population at time *t* = *t*_0_ by assigning each cell *i* an age *τ*_*i*_(*t*) at time *t* and molecule count vector ***x***_*i*_(*τ*_*i*_(*t*)).
2. *Intracellular reaction dynamics* .For each cell *i* simulate the trajectory of molecule counts in the age interval [*τ*_*i*_(*t*), *τ*_*i*_(*t* + ∆*t*)] using the Gillespie algorithm.
3. *Cell division times* For each cell *i* sample the division time *t*_*d,i*_ via the *thinning method*.
  a. Compute an upper bound of the division rate by *γ*_*max*_ ≥ *γ*(***x***(*τ* (*t*)), *τ* (*t*)) for all *t* in [*t, t*+∆*t*].
  b. Sample an event time *t*^*^ from exponential distribution with rate *γ*_*max*_. The proposed division time is then *t*_*p*_ = *t* + *t*^*^.
    - If *t*_*p*_ > *t* + ∆*t* then reject the proposed division time *t*_*p*_ and conclude that the cell does not divide in this time interval.
    - If *t*_*p*_ ≤ *t* + ∆*t*, sample a *u* from uniform distribution on the interval [0, 1] and consider two options. Accept the proposed time *t*_*p*_ as the next division time for the cell if 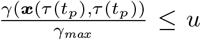. Otherwise, set *t* = *t*_*p*_ and repeat from step 3b.
4. *Next division time*. Determine the next cell to divide by *j* = argmin_i_*t*_*d,i*_.
5. *Cell division*. Replace the dividing cell *j* by two daughter cells at age 0. Set the molecule counts of one by sampling from the partitioning distribution *K*_1_(***x***|***x***_*j*_(*τ* (*t*_*d,j*_))) and give the remaining molecules to the other daughter. Simulate the intracellular reaction dynamics of the two daughter cells for the time interval [*t*_*d,j*_, *t* +∆*t*] and sample their division times as in Step 3.
6. Repeat the Steps 4-5 till no more divisions occur in the time interval [*t, t* + ∆*t*]. Set *t* = *t* + ∆*t* and ***x***_*i*_ to the molecule counts of cell *i* at time *t*. If *t* ≥ *T* stop, else go to Step 2.

### Finite state projection-based solution to the agent-based model

We developed the FSP scheme for growing cell populations. FSP relies on considering the dynamics over a finite truncated state space 𝒳 ⊂ 𝒮 for ***x***. The scheme effectively computes 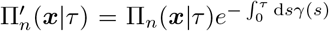, the product of the probability that cells have ***x*** molecules when they reach age *τ* and the probability that cells have not yet divided at that age. The cells that leave the truncated state space or do not divide in the age interval [0, *τ*_max_] are reinitialised at age 0 according to the division kernel *K*(***x***|***x***′). The corresponding exit probability to be reinitialised is given by

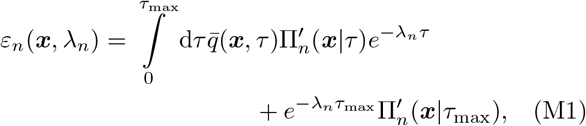

which can be self-consistently computed along with the FSP approximation. 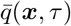 is the rate with which the cells with intracellular state ***x*** leave the truncated state space. That is, we define 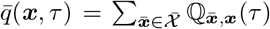 and the complement 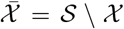 are the intracellular states outside the truncation boundary. The first term of (M1) corresponds to the probability mass leaving the population though the FSP boundary states into the complement 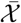 while second to the probability mass of cells that do not divide in the interval [0, *τ*_max_]. The algorithm is as follows:

1. Set *n* = 0 and provide an initial guess for the birth distribution 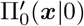 and growth rate *λ*_0_.
2. Solve

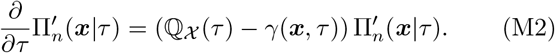

where ℚ _𝒳_ is defined as

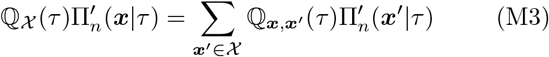

for all ***x*** in 𝒳.
3. Solve

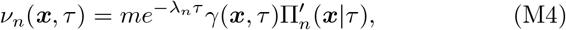

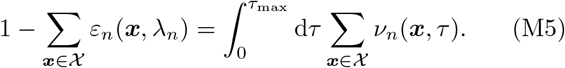

to obtain the growth rate *λ*_*n*_ and the division distribution *ν*_*n*_(***x***, *τ*)
4. Update the boundary condition 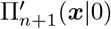 using

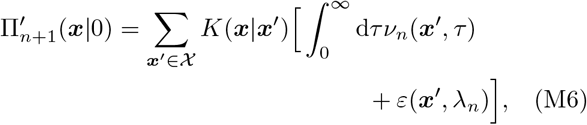
5. Repeat from 2. until *λ*_*n*_ and 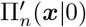converge.

For mother machine lineages, we have *m* = 1 and the population stays constant and hence *λ*_*n*_ = 0 for all *n*. In this case only the initial condition 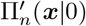 is iter-ated. The details of the derivation are given in SM Section S1 A.

### Modelling growth rate-dependent selection via effective selection strength

We consider an extended model where selection is modelled by gene expression-dependent growth and division rates. Cells grow exponentially where the cell growth rate *α*(***x***) is a continuous function of the gene expression state ***x***. The division rate *γ*(***x***, *ς, τ*) here also depends on cell size *ς*, which implements common models of cell size control such as adders and sizers. The stable snapshot distribution satisfies (SM Section S1 A; see also [60])

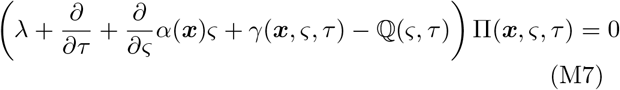

with boundary condition

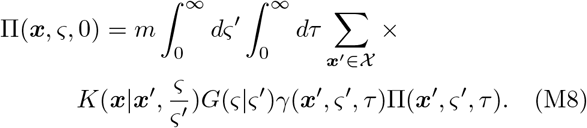

Here, the inherited size fraction *θ* has distribution *ρ*(*θ*), which influences the molecule-partitioning kernel *K*(***x***|***x***′, *θ*) and also determines size-partitioning via 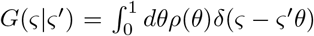 where *δ* is the Dirac delta function. This model reduces to Eqs. 1a to 1c by defining

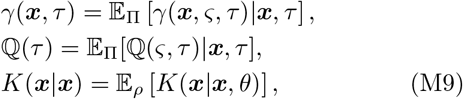

as the effective division rate, the effective transition matrix, and the effective partitioning kernel, respectively (SM Section S1 A). Note that, for cell size-dependent propensities, this amounts to using averaged propensities *w*_*r*_(***x***, *τ*) = 𝔼 _Π_[*w*_*r*_(***x***, *ς, τ*)|***x***, *τ* ] in (2).

To analyse dependence the effective division rate, we servations of the division distribution *ν*(***x***, *τ*) as state that the division rate can be written as

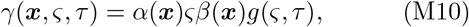

where now explicitly *α*(***x***) models growth rate-dependent selection and *β*(***x***) models division-rate selection and *g*(*ς, τ*) is the cell size control [56, 61]. This leads to:

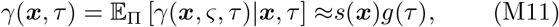

when neglecting the dependence of gene expression on cell size, i.e., 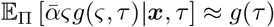and 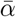 denotes an arbitrary reference growth rate without selection. Hence, growth rate-dependent selection can be modelled via an effective selection strength 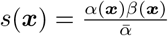.

### Inference from single-cell data

To perform parameter fitting for the agent-based models, we defined a likelihood based on *N* independent ob-servations of division distribution *ν*(*x, τ*) as

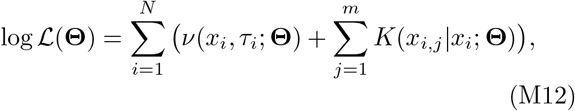

where 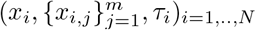 are the observations of protein counts of mother cell at division, protein counts of the daughter cells, and interdivision times, and where Θ is the vector of parameters of the model.

Inference of reaction kinetics and division rate requires absolute quantification of protein numbers. Assuming that the partitioning of molecules between the daughter cells is binomial with parameter 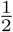, a linear relationship *f*_*i*_ = *a* · *x*_*i*_ between fluorescence *f* and absolute protein number *x*_*i*_ can be obtained (SM Section S4 A). In practice, one fits the relation between the partitioning variance of daughter fluorescence conditional on mother fluorescence (SM Fig. S8 and S10). The inference problem then becomes

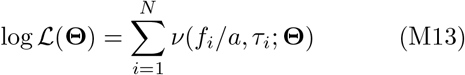

(*f*_*i*_, *τ*_*i*_)_*i*=1,..,*N*_ are the measured fluorescence intensity at division and the corresponding inter-division times. The details on the implementation are given in SM S4 C.

## Supplementary Materials

### S1. SNAPSHOT DISTRIBUTIONS IN AGENT-BASED POPULATIONS

We consider the biochemical reactions of the form

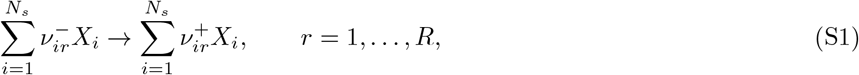

whose dynamics are encoded in the transition matrix:

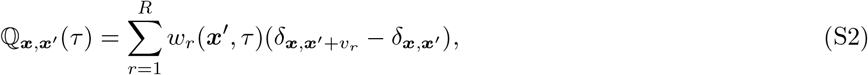

where *δ* is the Kronecker delta, *w*_*r*_(***x***′, *τ*) are the reaction propensities, and 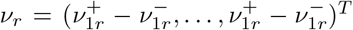 is the reaction stoichiometry. In the following we omit the age-dependence of the elements of the matrix ℚ(*τ*) and simply write *q*(***x, x***′) = ℚ_***x***,***x′***_ (*τ*).

As done in [22, 23] we derive the time-evolution of snapshot density *n*(***x***, *τ, t*) corresponding to the mean number of cells with molecule counts ***x*** and age *τ* at time *t* as

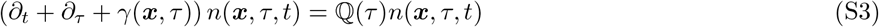

with the boundary condition

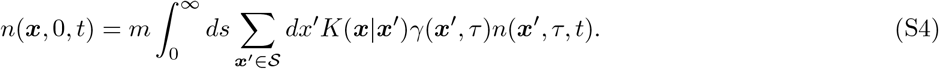

The sum is taken over the countable state space 𝒮 for the trait vector ***x***. The boundary condition corresponds to the cell divisions replacing a mother cell by *m* newborn daughter cells with *K*(***x***|***x***′) denoting the probability that a daughter cell inherits ***x*** molecules from a total of ***x***′ molecules of the mother cell. The division rate *γ*(***x***, *τ*) depends on both the cell age as well as the cell trait vector ***x***. Letting *n*(***x***, *τ, t*) ∼ *e*^*λt*^Π(***x***|*τ*)Π(*τ*), we obtain Eqs. (1). Finally *m* = 2 corresponds to the population lineage trees where each division results in two offspring while *m* = 1 corresponds to tracking a single offspring in mother machine lineages.

#### A. Finite State Projection method derivation

In this section we detail the construction of the finite state projection-based method for computing approximate solutions to the agent-based model. The method allows for the approximation error resulting from the truncation of the state space of the intracellular dynamics as well as error resulting from the finite time-horizon to be tracked and controlled. To that end, consider a truncated state space 𝒳 ⊆ 𝒮 for the trait ***x*** and let 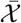 denote the complement of 𝒳. The evolution equations that consider only the evolution of cells that remain within the truncated state space 𝒳 is given by the following.

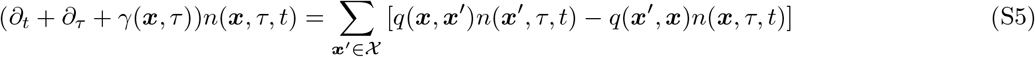

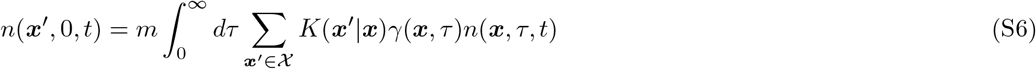

Let us denote

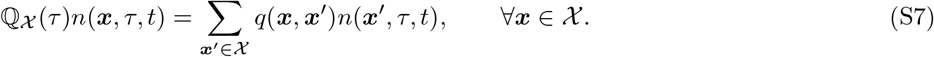

In addition, we keep track of the number of cells that exit the truncated state space 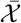 via the following evolution equation:

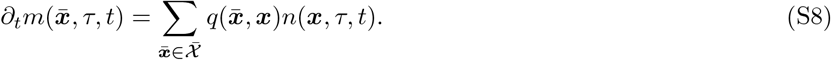

For brevity let us denote 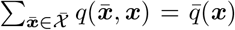. Note that the sum is taken over a potentially infinite complement 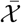. However, in practice when constructing the models based on a defined stochastic reaction network we can often easily find the total rate *q*(***x***) corresponding to reactions leaving the state ***x*** ∈ 𝒳. Further, we can also compute the total rate ∑_***x****′* ∈ 𝒳_ *q*(***x***′, ***x***) of transitioning from ***x*** to another state within the truncation 𝒳. We then use that 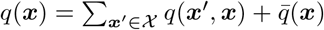 to evaluate the 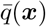 without having to sum over the infinite complement 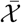.

To make analytical progress we consider the asymptotic behaviour of the defined system in the regime where the population grows exponentially with doubling rate *λ*. In particular, *n*(***x***, *τ, t*) ∼ *e*^*λt*^Π(***x***, *τ*) and 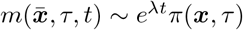 where Π and *π* are the snapshot densities of cells within the truncated state space and outside of it respectively. In the exponential growth regime the number of cells crossing the truncation boundary can then be given by

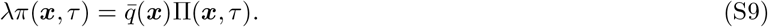

The additional source of error that needs to be considered is the finite time interval for integrals involving cell age. Let us fix a finite time interval [0, *τ*_max_]. The proposed method reinitialises both the cells that leave the truncation 𝒳 and cells that do not divide in the finite time interval [0, *τ*_max_] according to the division kernel as newborn cells.

Thus, we get the following system

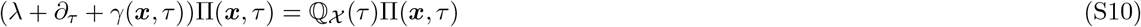

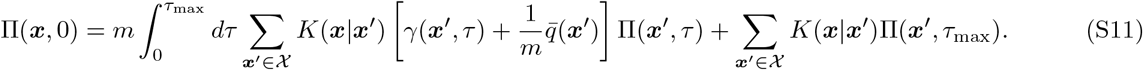

We then consider the conditional distribution 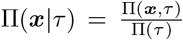. Substituting this into the joint evolution (Equation S10) and noting that the age-dependent marginal evolution is given by

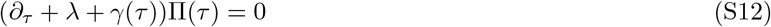

gives us

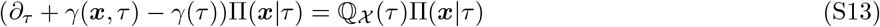

where *γ*(*τ*) is the marginal division rate 𝔼_Π_ [*γ*(***x***, *τ*)|*τ*] denoting the expectation taken with respect to Π(***x***|*τ*). We note that

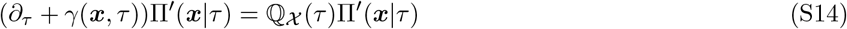

then describes the density of cells that have not divided upon reaching the age *τ* leading to the first passage time distribution for a cell to divide from trait ***x*** at time *τ*, division distribution for brevity, given by

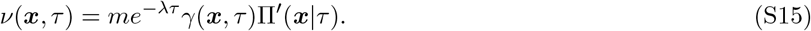

Again by the definition of conditional distribution we can then derive the boundary condition for the conditional evolution as

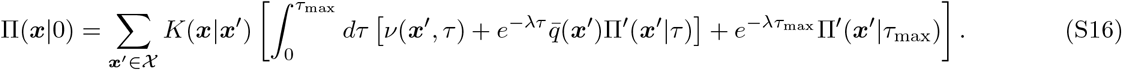

The exit probability

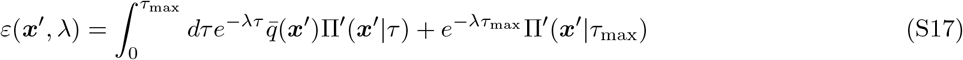

corresponds to the cells leaving the truncation from state ***x***′ and not dividing within the finite time horizon [0, *τ*_max_]. Note that summing over ***x***′ ∈ 𝒳 corresponds to the total exit probability *ε* for a cell to leave the state space

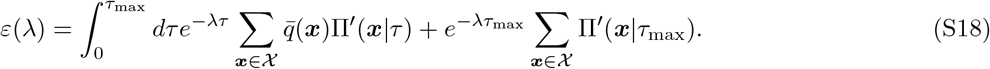

This takes into account truncation as well as the finite time-horizon used to compute the transient evolution of Π′(***x***|*τ*). Summing the boundary condition over ***x*** then gives

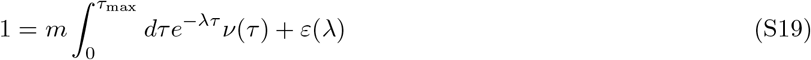

In the solution algorithm, presented in the Methods section of the main text, the equations (S16) and (S19) are computed at every iteration step. The implementation of the solution algorithm makes use of the Julia FiniteStateProjection package [62] for the construction of truncated ℚ(*τ*).

Note that if *ε*(***x***, *λ*) = 0 for all ***x*** ∈ 𝒳 then the boundary condition for the population lineage trees described in the main text by Equations (1) is recovered. Moreover

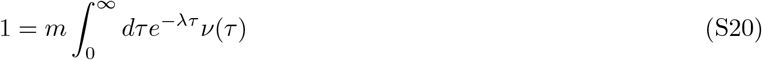

would correspond to the Euler-Lotka equation for the population lineage trees in the case of the untruncated state space and the time horizon [0, ∞].

### S2. RELATING GROWTH RATE SELECTION TO DIVISION RATE SELECTION

Here, we extend the snapshot density considered in Section S1 to cell size dynamics with growth-rate mediated selection. In particular, we assume that exponential growth of cell size *ς* with growth rate *α*(***x***), which continuously depends on the molecule numbers ***x***. We arrive at the following evolution equation for the snapshot density

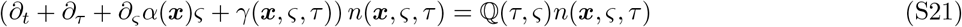

with boundary condition

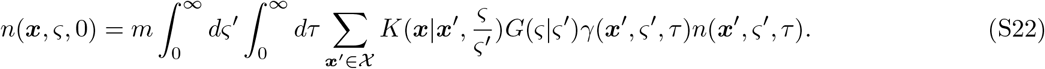

The partitioning of the molecule numbers is now assumed to depend also on the proportion of the size 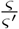 a daughter cell inherits from the mother cell of size *ς*′. The *G*(*ς*|*ς*′) defines the probability of a cell with size *ς*′ at division ending up with size *ς* after division. For example, for independent binomial partitioning, each molecule is partitioned into a daughter with a probability *θ* equal to the inherited size fraction, i.e.,

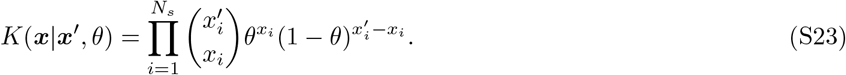

The inherited size fraction also defines the size division kernel:

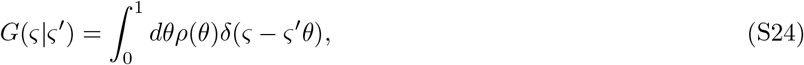

where *δ* is the Dirac delta function and *ρ* satisfies 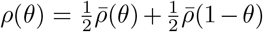 with 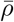 modelling the inherited size fraction accounting asymmetric cell division. The snapshot distribution in the asymptotic limit *t* → ∞ where the population grows exponentially with growth rate *λ* is then given by

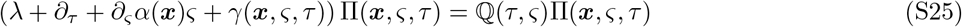

with boundary condition

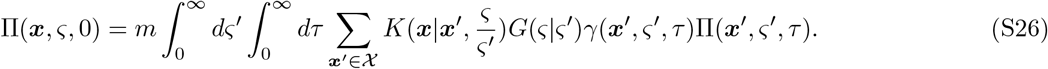

Integrating the division kernel over the birth sizes *ς*

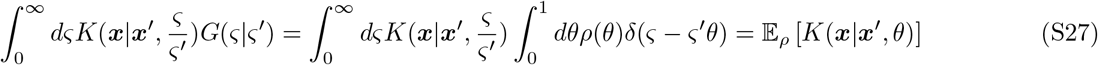

Using the law of conditional probability Π(***x***, *ς, τ*) = Π(*ς*|***x***, *τ*)Π(***x***, *τ*) and marginalising out *ς*, the evolution equation for the marginal snapshot distribution with boundary condition becomes

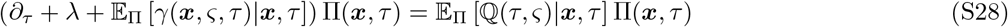

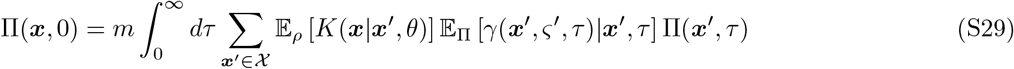

We define

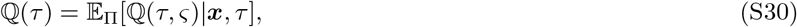

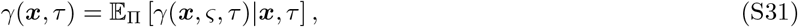

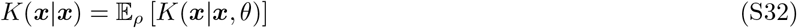

as the marginal division rate and the effective division kernel, respectively. We then recover the model that depends only on the gene expression state ***x*** and the cell age *τ* :

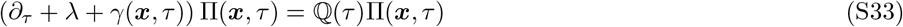

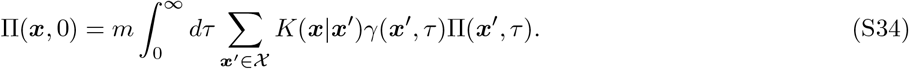

The model without cell size control is thus a reduced model of the developed cell size control model.

### S3. MODEL DESCRIPTIONS

#### A. Telegraph model

In this section we provide the complete specification of the agent-based telegraph model used in the main manuscript. The intracellular dynamics are given by the following stochastic reaction network

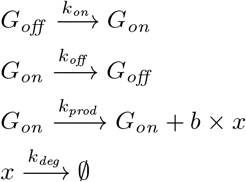

where 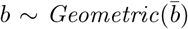 is a geometrically distributed random variable parametrised by the mean 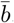. In practice, to simplify the implementation with Julia Catalyst package [63] the bursty reaction was truncated and split into multiple reactions

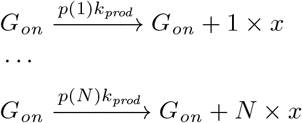

where the rate of each reaction is weighed by the probability density *p* of the burst size. Note that the truncation of the bursty reaction is not necessary for the computations of 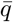 in Section S1 A. The total rate out of the each state with gene on and protein count *x* is the sum of *k*_*off*_, *k*_*deg*_ and *k*_*prod*_. Thus all we need to find the rate corresponding to cells leaving the truncation is to consider the cumulative probability of bursts that do not leave the truncation.

In the case of neutral condition with no selection the division rate is given by *γ*(***x***, *τ*) = *g*(*τ*) where *g*(*τ*) is the hazard of the Gamma distribution parametrised by mean its *μ* and squared coefficient of variation *cv* ^2^. In general, if *f* and *F* are the probability density and cumulative distribution functions of a distribution respectively, then the hazard of the distribution is defined by

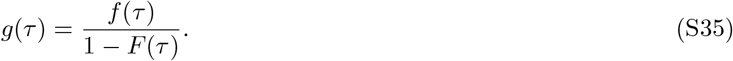

In the selection case the division rate is given by *γ*(***x***, *τ*) = *s*(***x***)*g*(*τ*) where *s*(*x*) is a Hill function

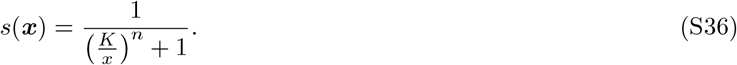

The parameters are chosen to describe the slow-switching dynamics of the promoter state to give rise to bimodal expression levels and are given in the table below.

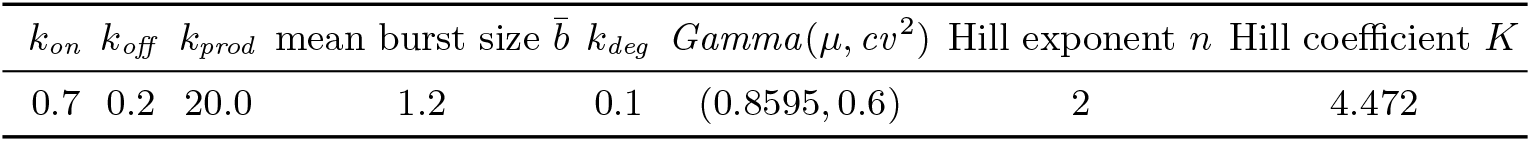

#### B. Single gene feedback models

There are several models that follow the same general structure presented in the main text. We start by presenting the general structure of the agent-based and effective dilution formulations of the single gene feedback models. The specific instances of these models are then arrived to by choosing different ways to define the rate functions within the models.

##### 1. Agent-based model

The intracellular dynamics of the agent-based model are given by the following general stochastic reaction network

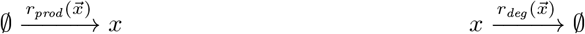

modelling production and degradation of a single protein *x* with rates 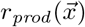 and 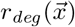 respectively. The general model assumes the division rate *γ*(***x***, *τ*) is in the form

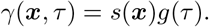

The partitioning of mother cell protein numbers at division follows symmetric binomial partitioning.

##### 2. Effective dilution model

In the effective dilution model we add a linear dilution term to the stochastic reaction network

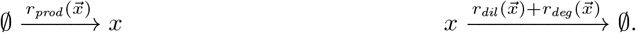

where 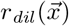 is the rate of dilution modelling the loss of protein due to cell divisions in an exponentially growing population. A common approach when no division-rate selection on *x* is considered is to take 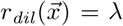 where *λ* is the doubling time of the cell population. The stationary solutions for the birth-death processes arising from the effective dilution models were computed using standard steady state methods [64].

##### 3. Model instances

There are three different variants of the single protein feedback model that are considered in this paper. Each of them use the same general outline defined in Sections S3 B 1 and S3 B 2.

###### a. Transcriptional feedback model

The transcriptional feedback model considers the protein *x* promoting its own production without division-rate selection on *x* by defining the rate of production and degradation as

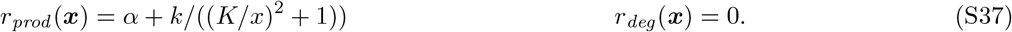

The degradation rate of the protein is taken to be 0 for simplicity. This model considers no division-rate selection on *x* and thus we take *s*(***x***) = 1. The age-dependent division-rate component *g*(*τ*) is given as the hazard of the Gamma distribution parametrised by its mean *μ* and the squared coefficient of variation *cv* ^2^. Let *f* and *F* denote the probability density and cumulative distribution functions of the Gamma distribution respectively. The division rate *γ*(***x***, *τ*) is then given by

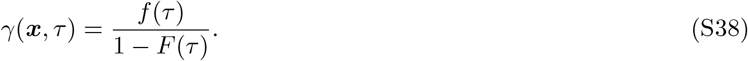

The dilution rate in the effective dilution model is defined as

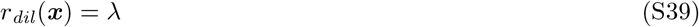

where *λ* is the doubling time of an exponentially growing cell population resulting from the Gamma distributed interdivision time distribution above. The parametrisations chosen to give rise to bimodal behaviour are given in the table below.

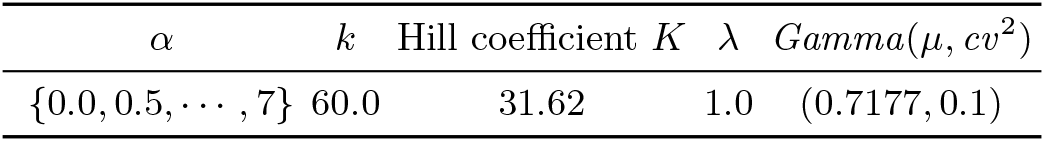

###### b. Positive growth feedback model

The positive growth feedback model uses the general structure outline in Sections S3 B 1 and S3 B 2 and features a constant production and degradation rate

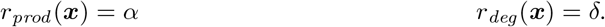

The *x*-dependent division rate function *s*(***x***) is given by a repressive Hill function

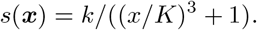

while the age-dependent component is taken to be constant *g*(*τ*) = 1.0 corresponding to exponential division time distribution when no selection on gene expression is considered. The dilution rate in the effective d ilution m odel is defined as

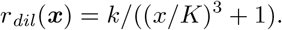

The parametrisations of the model are given in the table below.

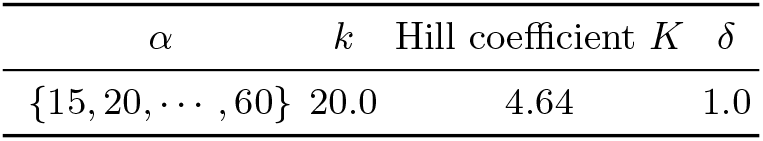

###### c. Combined feedback model

The combined feedback model combines the transcriptional feedback mechanism from Section S3 B 3 a and growth feedback mechanism from Section S3 B 3 b. In particular,

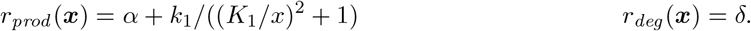

The division rate function is similar to growth feedback model (with higher Hill exponent) and is given by a Hill function dependent on the protein counts

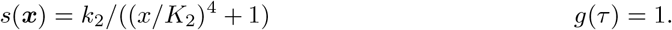

In the effective dilution model consider the corresponding dilution term

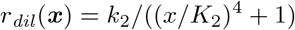

The parametrisations chosen to give rise to bimodal behaviour are given in the table below.

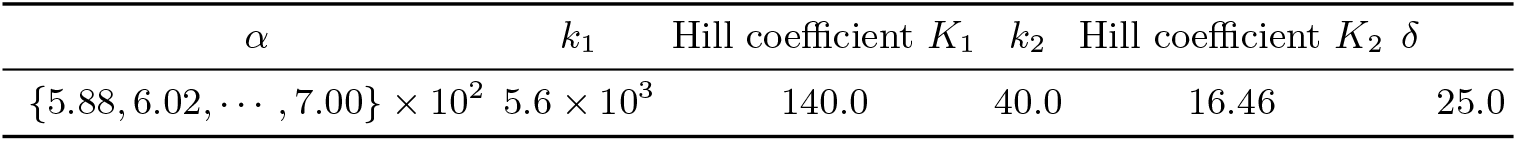

#### C. Genetic toggle switch

Here we provide the complete specifications of the genetic toggle switch model used in the main manuscript.

##### 1. Agent-based model

The intracellular dynamics of the agent-based model are given by the following stochastic reaction network

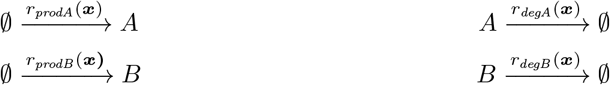

where the functions defining the rates of production and degradation for the two protein counts *x*_*A*_ and *x*_*B*_ are as follows:

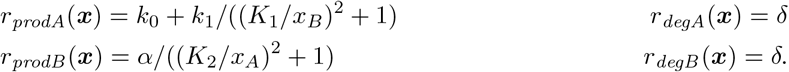

In particular, the proteins *A* and *B* inhibit each other’s expression. We call *α* the induction strength. The division rate *γ*(***x***, *τ*) for the agent-based model is then given by

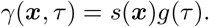

Where

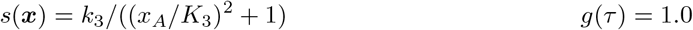

In particular, we are considering positive division-rate selection on the protein *A*.

##### 2. Effective dilution model

As previously, in the effective dilution model we add a linear dilution term to the stochastic reaction network

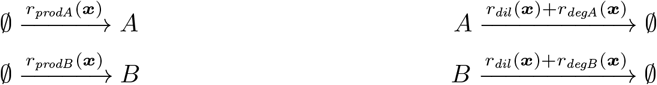

where *r*_*dil*_ (***x***) is the rate of dilution modelling the loss of protein due to cell divisions in an exponentially growing population. The dilution rate in the effective dilution model is defined as

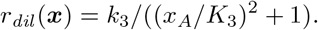

The parametrisations of the model are given in the table below.

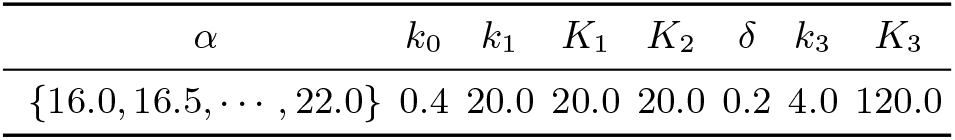

### S4. PARAMETER AND DIVISION RATE INFERENCE

#### A. Conversion factor

The *Escherichia coli* datasets [36, 47] analysed in the main text report fluorescence intensity of the reporter protein. In order to fit the agent-based model to data we convert the intensity to copy numbers of the protein by analysing the variability in the mother and daughter cell fluorescences at division as done, for example, in [65]. To that end we find a linear relationship *f* = *ax* between the copy numbers of fluorescent molecules *x* and fluorescence intensity *f*. This conversion is done under the assumption that partitioning of the fluorescent molecules between the two daughter cells at division is binomial with probability 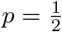. Variance of the daughter cell fluorescence *f*_*d*_ is then given by

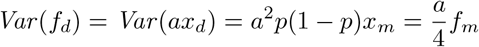

where *x*_*m*_ and *f*_*m*_ are the molecule count and fluorescence of the mother cell respectively. From the data we then fit the slope *ā* of *Var* (*f*_*d*_) = *āf*_*m*_ and equate it with 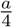 to estimate the conversion factor *a*.

#### B. Bursty gene expression model

The agent-based model considered for both the DNA damage response and antibiotic resistance data sets is the same with different parametrisations. The intracellular dynamics of the agent-based model are given by the following stochastic reaction network

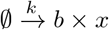

where 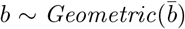 is a geometrically distributed random variable parametrised by the mean 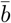 (see Section S3 A for implementation details). The division rate of the model is given by *γ*(***x***, *τ*) = *s*(***x***)*g*(*τ*) where *g*(*τ*) is the hazard of the kernel density estimate of the interdivision times in the data. If *f* and *F* denote the probability density and cumulative distribution functions of the kernel density estimate

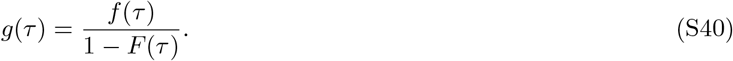

The function *s*(***x***) is fitted to individual conditions. In the cases of no antibiotic treatment and no induced DNA damage we assume there is no division-rate selection on the gene expression and thus the *s*(***x***) = 1. In the cases of antibiotic treatment and induced DNA damage we fit the parameters of a Hill function of the form

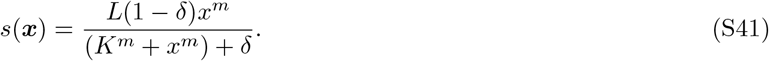

The partitioning of mother cell molecule numbers at division follows symmetric binomial partitioning. The parameters fitted for the DNA damage response model is given in Table S1 and the parameters fitted for the antibiotic resistance model are given in Table S2.

**TABLE S1.** The parameters fitted for the DNA damage response model via Bayesian Optimisation.

| | $k$ | $b$ | $L$ | $K$ | $\delta$ | $m$ |
| --- | --- | --- | --- | --- | --- | --- |
| Wild type | 0.2070 | 1.4643 | N/A | N/A | N/A | N/A |
| DNA damage | 0.2070 | 1.4643 | 1.0469 | 44.68 | 0.0 | 11 |

**TABLE S2.** The parameters fitted for the antibiotic resistance model via Bayesian Optimisation.

| | $k$ | $b$ | $L$ | $K$ | $\delta$ | $m$ |
| --- | --- | --- | --- | --- | --- | --- |
| No treatment | 0.5684 | 1.4087 | N/A | N/A | N/A | N/A |
| Treatment (selection) | 0.5684 | 1.4087 | 1.2336 | 70.4 | 0.0 | 20 |
| Treatment (selection + adaptation) | 0.3378 | 2.0725 | 1.4643 | 92.8 | 0.0 | 4 |

#### C. Bayesian optimization

The Bayesian optimisation routine uses the `scikit-optimize` Python package with the default options [66]. In particular, the optimiser uses the Matérn kernel with smoothness parameter *ν* = 2.5 corresponding to twice differentiable functions. The length scales of the kernel are tuned by the optimising routine by maximising the log marginal likelihood. Finally, the acquisition function is chosen probabilistically at every iteration between the lower confidence bound, negative expected improvement and negative probability improvement acquisition functions implemented in the package. We performed the parameter optimisation with 200 evaluations of the likelihood function and initialised with 10 random parametrisations. This was replicated 5 times with the optimal parametrisation across the replications chosen. The bounds used to constrain the search space are given in the table below.

| $k$ | $b$ | $L$ | $K$ | $\delta$ | $m$ |
| --- | --- | --- | --- | --- | --- |
| (0.0, 1.0) | (1.0, 10.0) | (0.0, 5.0) | (0.0, 150.0) | (0.0, 1.0) | (0, 20) |

**FIG. S1.**
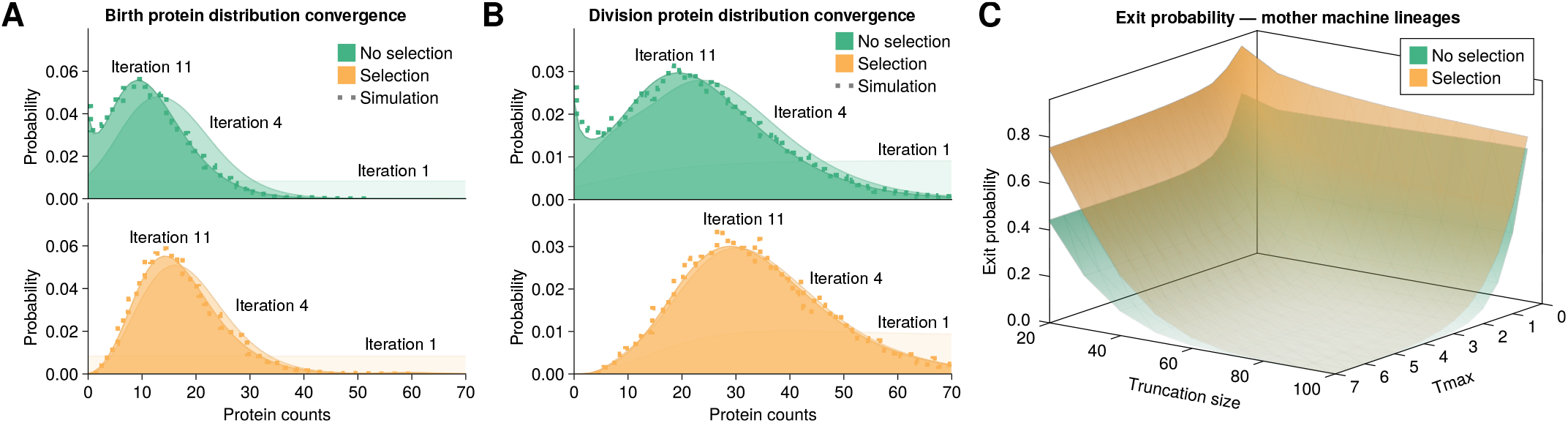
Convergence of the mother machine lineage statistics of the telegraph model. **(A-B)** Birth (panel **A**) and division (panel **B**) distributions of the mother machine lineage model with fixed a truncation size and *τ*_max_ converges in a couple of iterations and agrees with the agent-based simulation. **(C)** The exit probability due to the FSP truncation decreases with the truncation size and time horizon *τ*_max_.

**FIG. S2.**
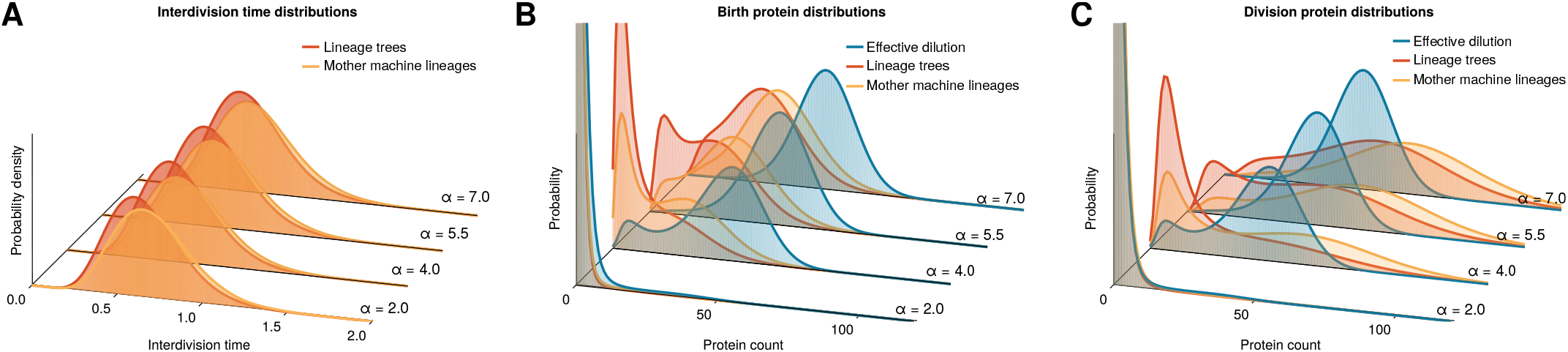
Summary distributions of the transcriptional feedback model. **(A)** The interdivision time distribution shows slow dividing cells being over-represented in mother machine lineages (yellow) compared to population lineage trees (red). **(B-C)** Protein distributions predicted by the agent-based models at birth (panel **B**) and division (panel **C**) are compared with the stationary distribution of the effective dilution model.

**FIG. S3.**
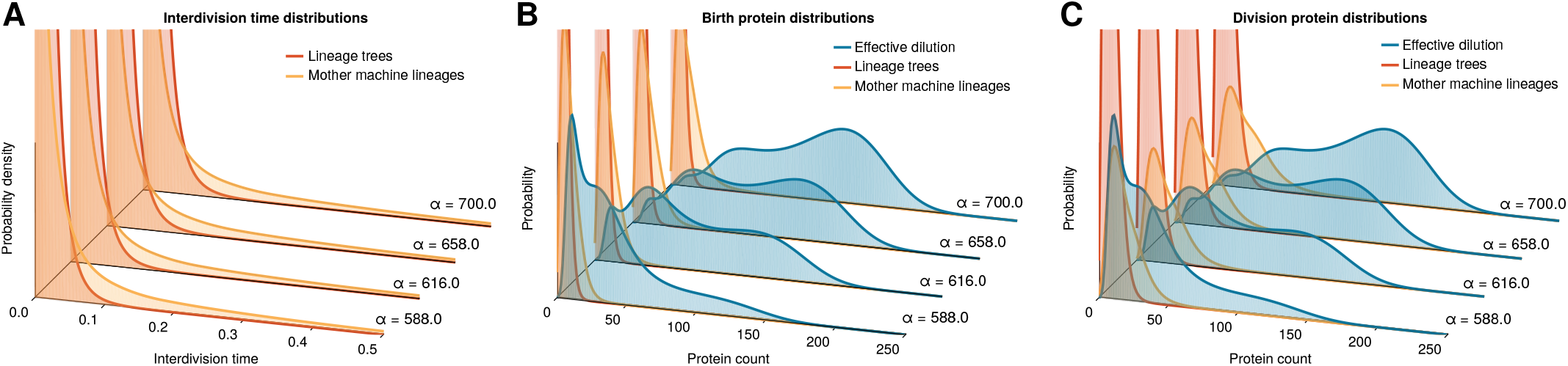
Summary distributions of the model combining transcriptional and growth feedback mechanism. **(A)** The interdivision time distributions display the behaviour observed for the growth feedback model with the right tail of the unimodal interdivision time distribution becoming longer as the synthesis rate *α* increases. As before the corresponding slow dividing cells are repressed in the lineage tree statistics. **(B-C)** Protein distributions predicted by the agent-based models at birth (panel **B**) and division (panel **C**) are compared with the stationary distribution of the effective dilution model.

**FIG. S4.**
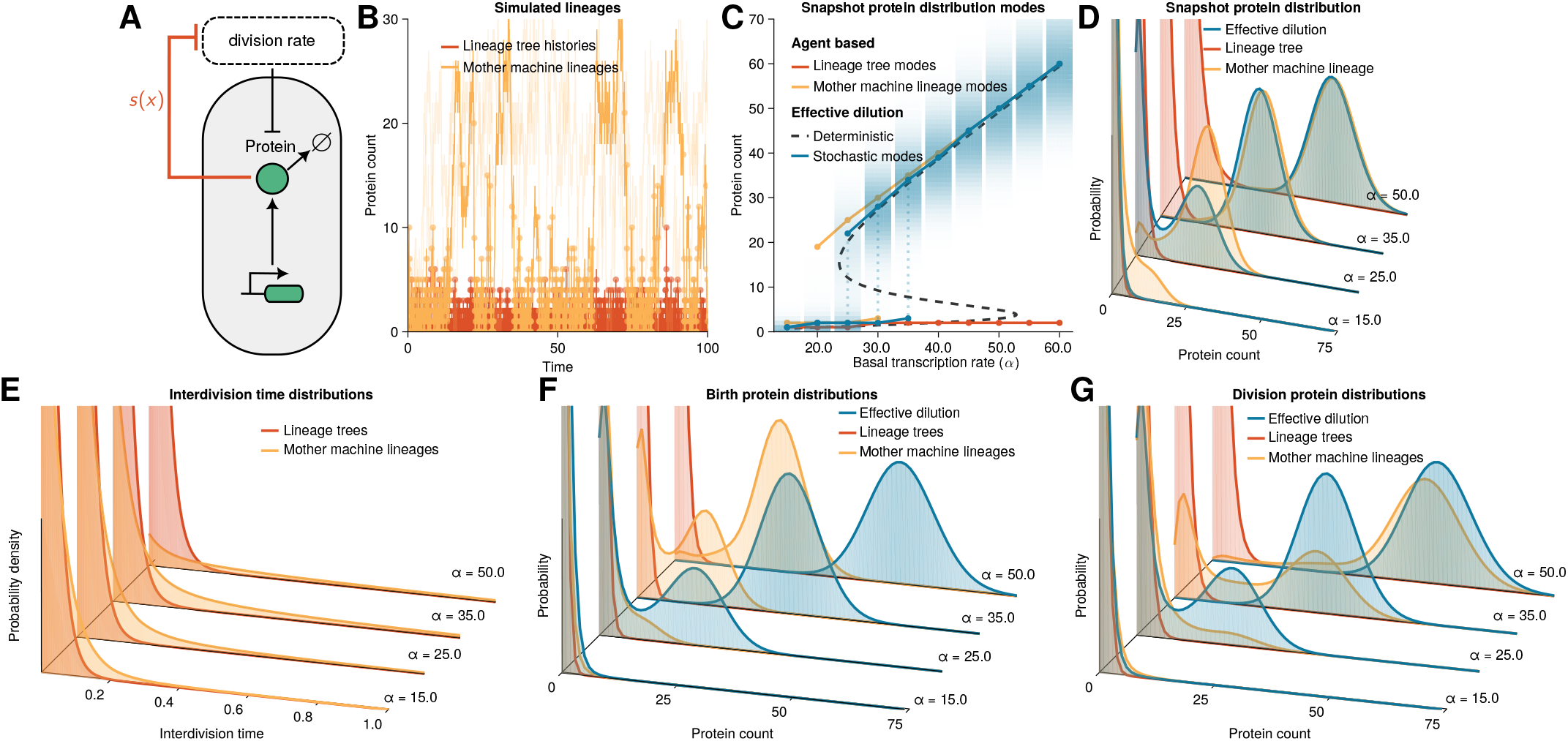
**(A-D)** Agent-based model with growth feedback (SI Section S3 B 3 b). **(A)** Illustration of the growth feedback model where stable proteins are synthesized with rate *α* and high protein abundance inhibits cell divisions. **(B)** Agent-based simulations of mother machine lineages (*α* = 20.0) show switching between low and high protein levels while, in lineage tree histories, the fast dividing lineages determine the cell fate. **(C)** Protein distribution modes display bimodality for mother machine lineages and EDM but only one mode for lineage trees. **(D)** Protein distribution of mother machine lineages display slowly diving subpopulations not seen in lineage trees. **(E)** The interdivision time distribution shows slow dividing cells are over-represented in mother machine lineages (yellow) compared to the population lineage trees (red). The right tail of the unimodal interdivision time distribution corresponding to the mother machine lineages becomes longer as the protein production rate *α* increases. The corresponding slow dividing cells are repressed in the lineage tree statistics. **(F-G)** Protein distributions predicted by the agent-based models at birth (panel **F**) and division (panel **G**) are compared with the stationary distribution of the effective dilution model.

**FIG. S5.**
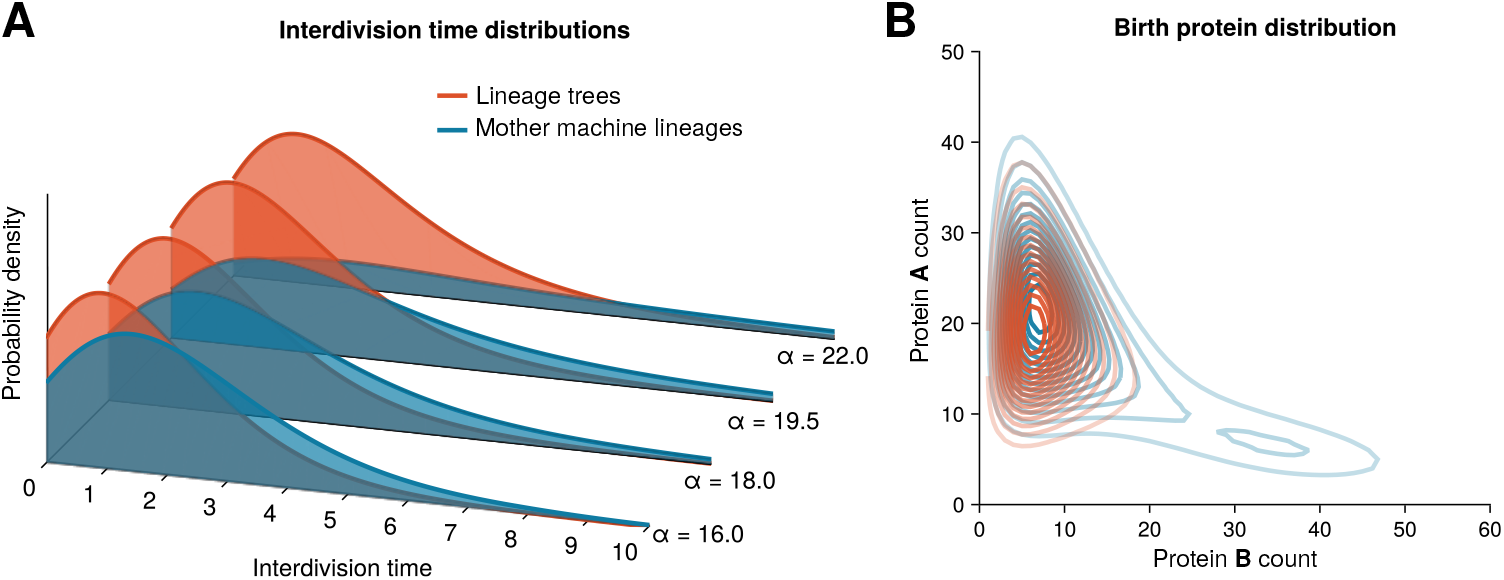
Summary distributions of the genetic toggle switch (SI S3 C). **(A)** The right tail of the unimodal interdivision time distributions becomes longer as the synthesis rate *α* increases. The corresponding slow dividing cells are repressed in the lineage tree statistics compared to the mother machine lineages. **(B)** Birth protein distributions predicted by the agent-based models (*α* = 18.9) for mother machine lineages (blue) displays bimodality with the less prominent peak corresponding to slow-dividing cells while in lineage trees (red) this peak is missing.

**FIG. S6.**
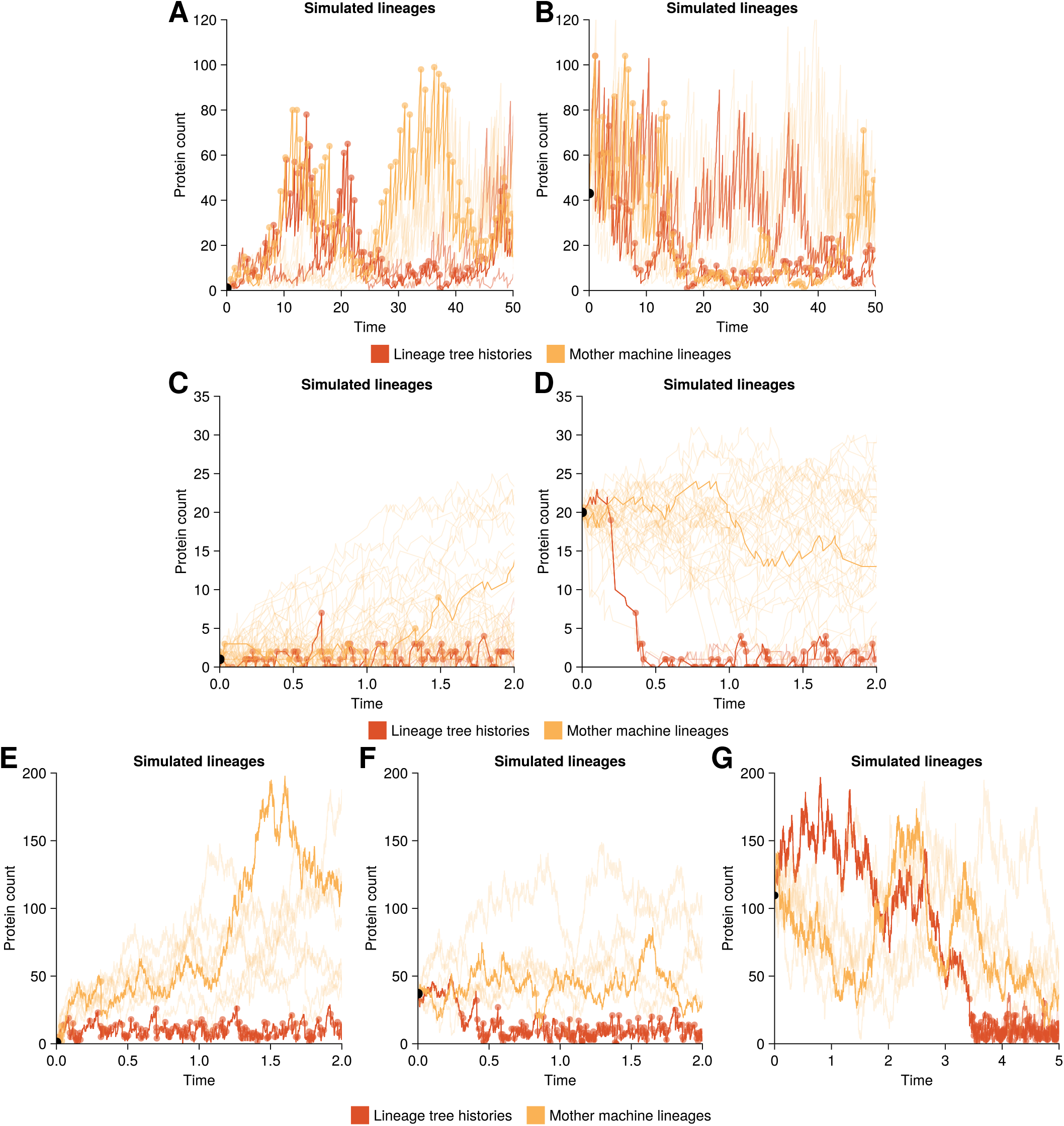
Agent-based stochastic simulations of lineage tree histories (red) and mother cell lineages (yellow) of the single gene feedback models. **(A-B)** Simulations of transcriptional feedback model (model details in SI Section S3 B 3 a) for *α* = 4.0. Agent-based simulations with cells initialised in a low expression level state (1 molecule, panel **A**) and a high expression level state (20 molecule, panel **B**). Lineage tree histories and mother machine lineages explore both the high expression and low expression states under both initialisations. **(C-D)** Simulations of the growth feedback model (model details in SI Section S3 B 3 b) for *α* = 20.0. **(C)** Agent-based simulations initialised with cells in a low expression level fast dividing state (1 molecule). The lineage tree histories remain in their initial fast dividing state while mother machine lineages eventually visit the slow dividing state. **(D)** Agent-based simulations initialised with cells in a high expression level state (20 molecules). Lineage tree histories show strong selection for the fast dividing cell lineages in a population with histories coming down to the low expression fast dividing state in a couple of divisions. The histories resulting from cells that quickly make the transition to fast dividing regime are going to be highly represented in lineage tree histories. **(E-G)** Simulations of the combined feedback model (model details in SI Section S3 B 3 c) for *α* = 616.0. Agent-based simulations with cells initialised in a low expression level state (1 molecule, panel **E**) and two high expression level states (37 molecules in panel **F**, 111 molecules in panel **G**). On short time scales the mother machine lineages and lineage tree histories agree with the qualitative dynamics of each other. However, over time the lineage tree histories settle in the fast dividing cell state.

**FIG. S7.**
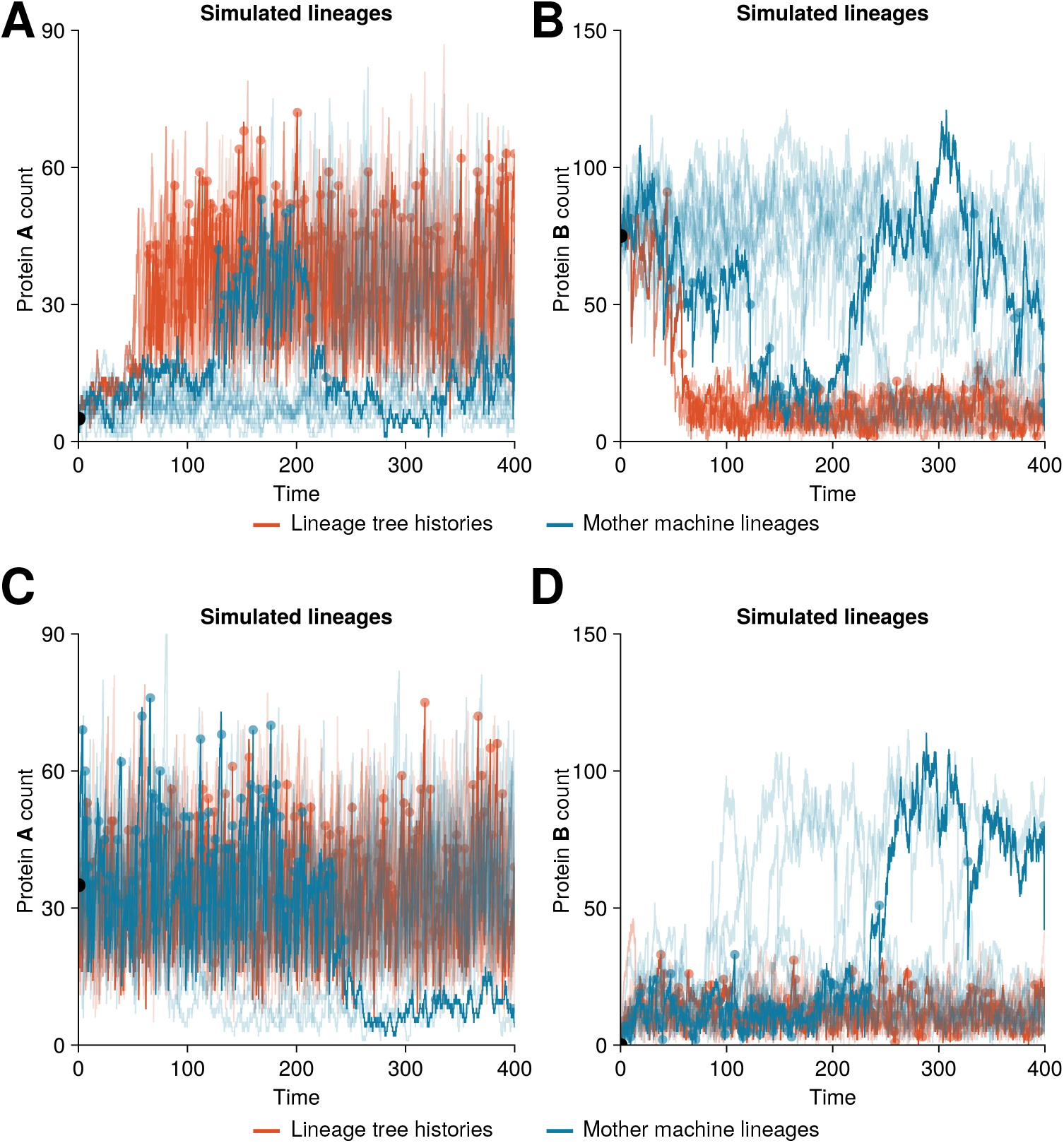
Agent-based stochastic simulations of lineage tree histories (red) and mother cell lineages (blue) of the genetic toggle switch for induction strength *α* = 18.9 (model details in SI S3 C). **(A-B)** Simulations trajectories starting with a cell in a slow dividing state (protein A count 5, B count 75). **(C-D)** Simulations trajectories starting with a cell in a fast dividing state (protein A count 35, B count 0).

**FIG. S8.**
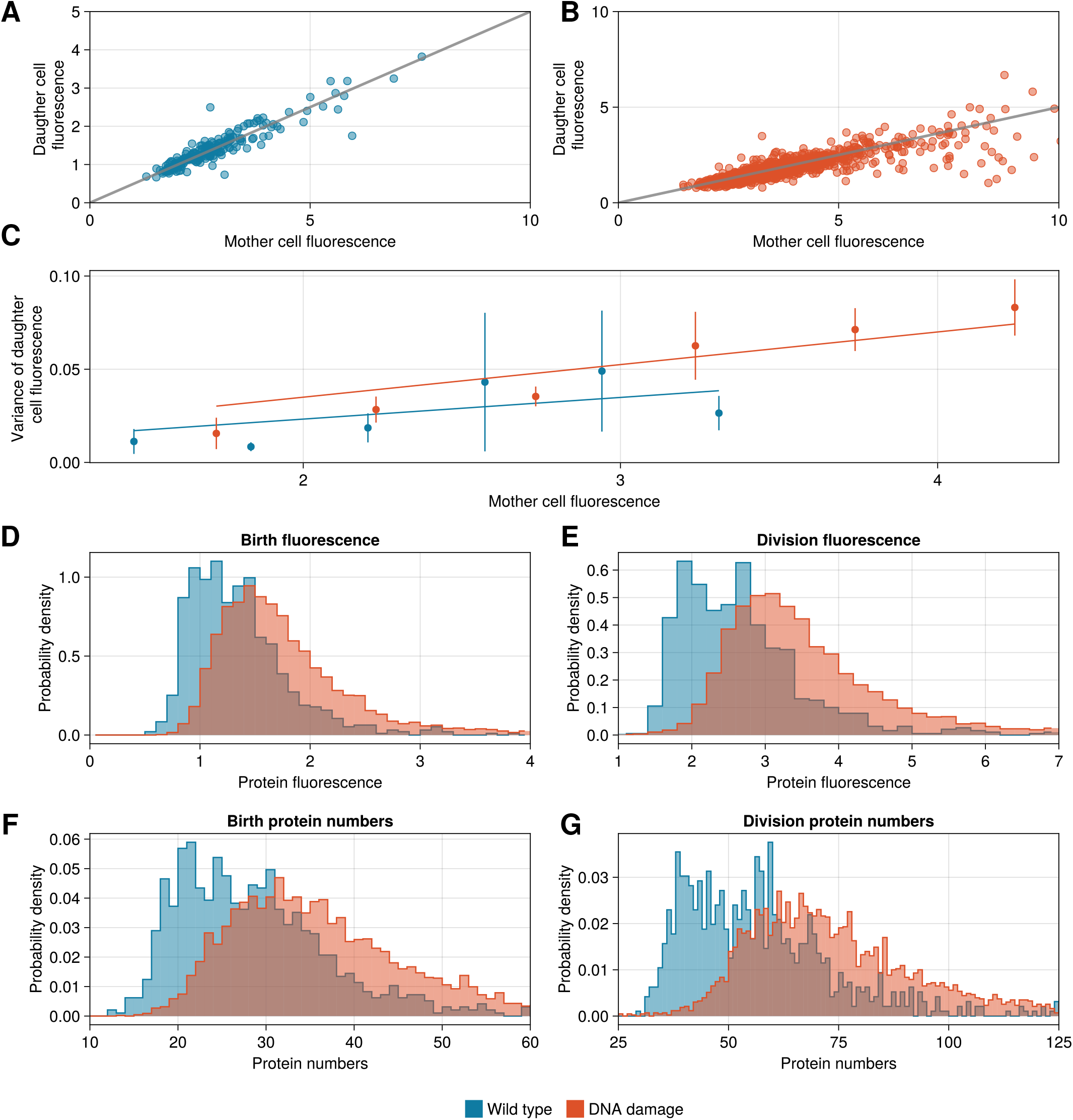
Conversion factor estimation for the DNA damage response dataset. **(A-B)** The partitioning of fluorescence between the mother and daughter cells compared against symmetric partitioning (gray line with slope 0.5). **(C)** Linear regression lines for the binned mother cell fluorescence values against the variance of the daughter cell fluorescence within the bin. Scatter points represent the midpoints of the bins. The slope of the wild type case is chosen as the conversion factor. **(D-E)** Histograms displaying the birth (panel **D**) and division (panel **e**) fluorescence distributions of the wild type (blue) and the induced DNA damage strain (red) from the data. **(F-G)** Histograms showing the birth (panel **F**) and division (panel **G**) protein distribution resulting from the estimated conversion factor.

**FIG. S9.**
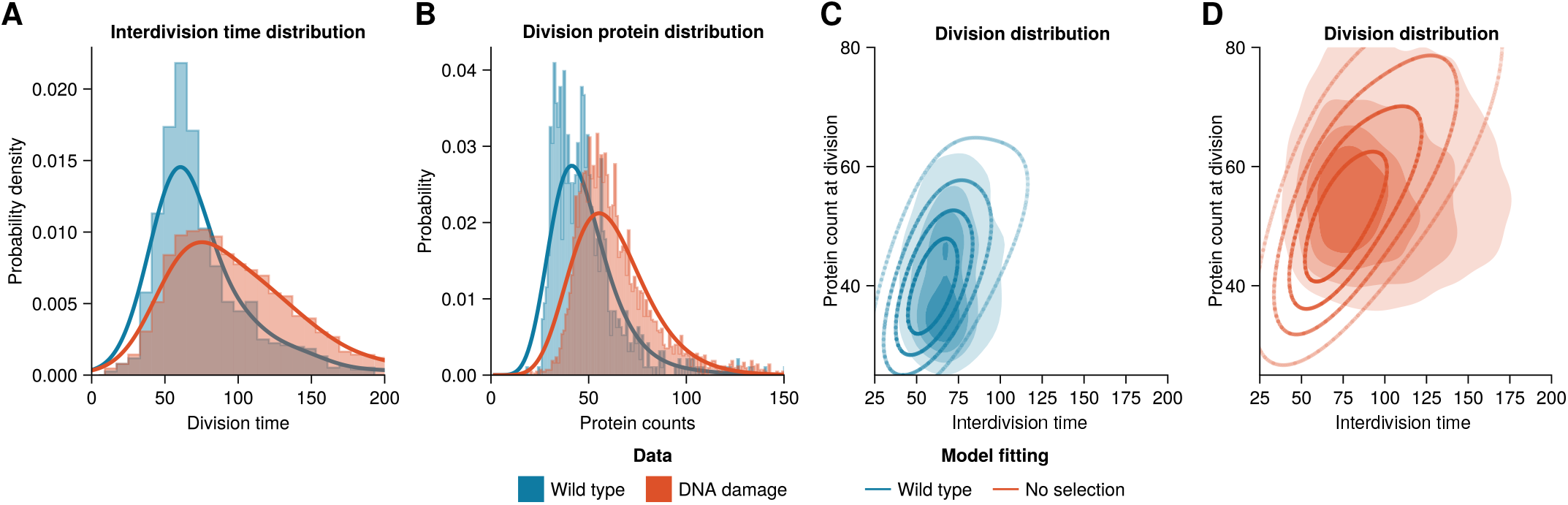
Changes in the division rate without selection effects do not capture the SOS response dynamics observed in the data. **(A)** Interdivision time distributions for the two conditions are fitted by a separate kernel density estimate. **(B)** With the gene expression dynamics kept constant between the conditions the division protein distribution is well captured by the model. **(C-D)** Comparison of the division distributions of the wild type (panel **C**) and damage-induced (panel **D**) cells from the data with the mother machine lineage division distribution. The model overestimates the correlation between the protein count and interdivision time for the damage-induced cells.

**FIG. S10.**
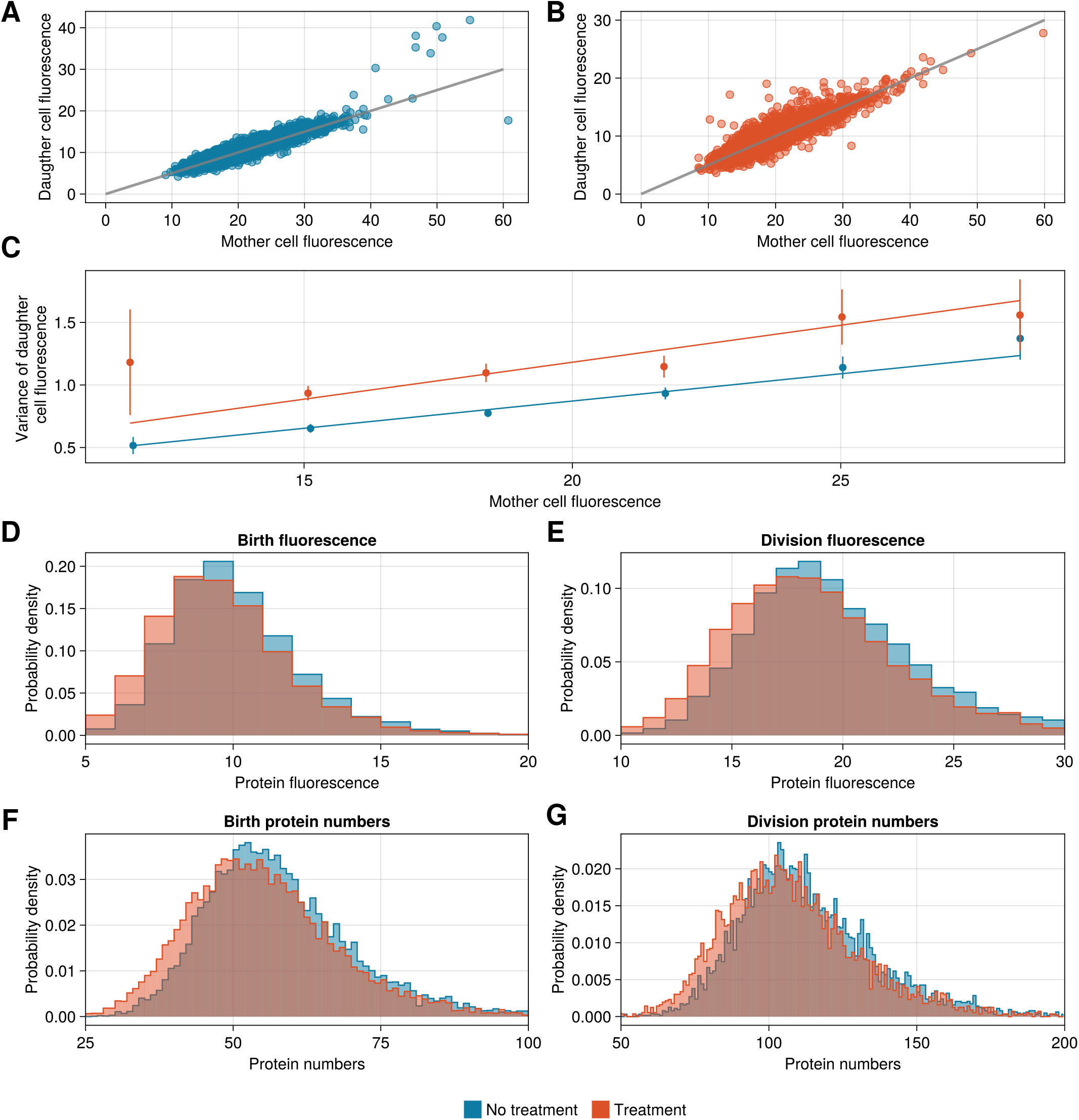
Conversion factor estimation for the DNA damage response dataset. **(A-B)** The partitioning of fluorescence between the mother and daughter cells compared against symmetric partitioning (gray line with slope 0.5). **(C)** Linear regression lines for the binned mother cell fluorescence values against the variance of the daughter cell fluorescence within the bin. Scatter points represent the midpoints of the bins. The slope of the no treatment case is chosen as the conversion factor. **(D-E)** Histograms displaying the birth (panel **D**) and division (panel **E**) fluorescence distributions of the untreated (blue) and with antibiotic-treated (red) cells from the data. **(F-G)** Histograms showing the birth (panel **F**) and division (panel **G**) protein distribution resulting from the estimated conversion factor.

**FIG. S11.**
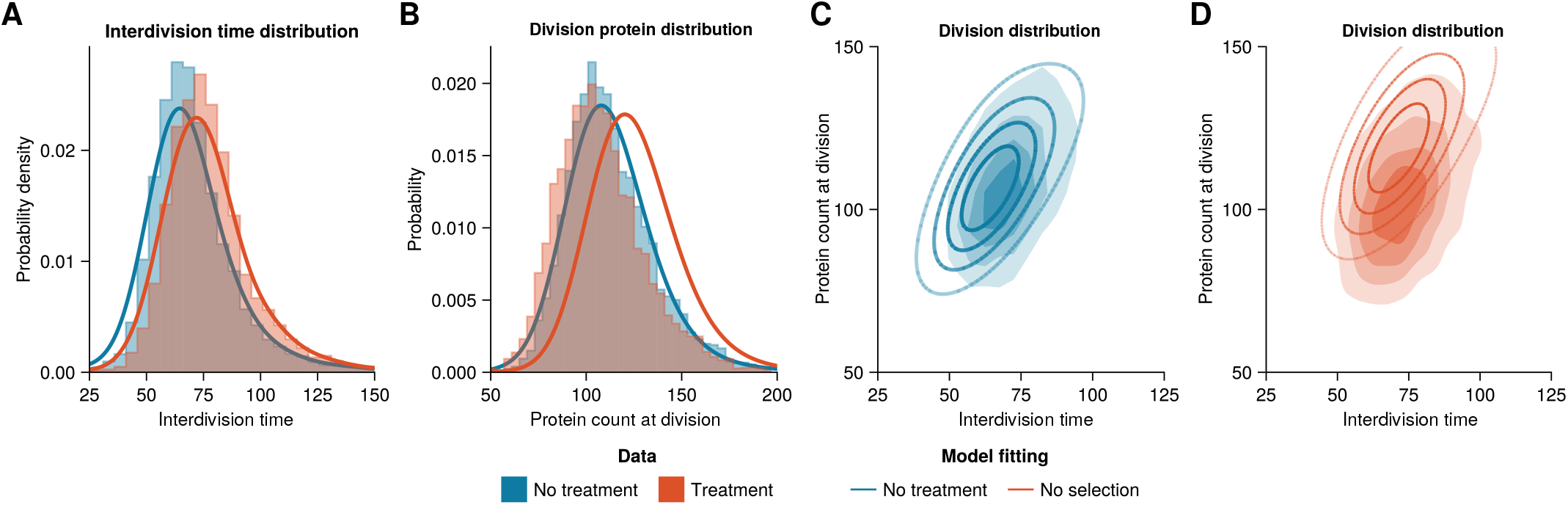
Changes in the division rate alone are not sufficient to capture the antibiotic treatment response of *Escherichia coli* cells. **(A)** Interdivision time distributions of for conditions are fitted by a separate kernel density estimate. **(B)** With the gene expression dynamics kept constant between the conditions the model predicts significantly higher division protein counts than observed in the antibiotic-treated data. **(C-D)** Comparison of the division distributions of untreated (panel **C**) and antibiotic-treated (panel **D**) cells from the data with the corresponding agent-based lineage tree division distributions.

